# Multiple genes evolved for fungal septal pore plugging identified via large-scale localization and functional screenings

**DOI:** 10.1101/2022.11.24.517796

**Authors:** Md. Abdulla Al Mamun, Wei Cao, Shugo Nakamura, Jun-ichi Maruyama

**Author notes:** Address correspondence to Jun-ichi Maruyama, Department of Biotechnology, The University of Tokyo, Tokyo, Japan.

## Abstract

Multicellular organisms exhibit cytoplasmic exchange using porous structures for cooperation among cells. Fungal multicellular lineages have evolved septal pores for this function. Interconnected hyphal cells possess the risk of wound-related cytoplasmic loss unless the septal pores are plugged. However, the gene evolution of regulatory mechanisms underlying fungal septal pore plugging remains poorly understood. To identify novel septal components, 776 uncharacterized proteins were identified using genomic comparisons between septal pore-bearing and -lacking ascomycete species. We then determined their subcellular localizations, and in total 62 proteins localized to the septum or septal pore. We analyzed the effects of deleting the encoding genes on septal pore plugging upon hyphal wounding. Of the 62 proteins, 23 were involved in regulating septal pore plugging. Here, using orthologous group and phylogenetic analyses, this study suggests that septal pore regulation has evolved either by co-option of preexisting genes or by Pezizomycotina-specific gene acquisition.

## INTRODUCTION

The emergence of multicellularity represents an important transition in evolutionary history. Organisms from diverse origins have organized into simple multicellular morphologies such as filaments, clusters, and sheets, where intercellular communication is limited due to a lack of direct passageways in the subsequent cell population^1,2^. In relatively late evolutionary history, several groups of eukaryotic organisms have developed complex multicellularity via cell-cell adhesion and developmental programs to differentiate into tissues, reproductive organs, and fruiting bodies^1,3^. In addition, complex multicellular organisms have evolved cell-to-cell connectivity via ultrastructural passageways to facilitate signal transmission^4^. Such cell-to-cell connectivity has independently evolved in animals, plants, and multicellular fungi; moreover, its regulation confers selective advantages in unfavorable environments. In animals, gap junctions, which allow ions and small molecules to pass, are perturbed structurally and functionally by oxidative stress^5,6^. In plants, plasmodesmata, which mediate cell-to-cell trafficking of transcription factors and signaling molecules, can be blocked by callose deposition in response to abiotic stresses^7,8^.

Fungi include both septal pore–bearing (filamentous fungi) and –lacking (yeast) species, facilitating the characterization of septal pore-related organization using comparisons. Fungal hyphae grow by extending polarized tips and further compartmentalize by septa in relation to cell size and nuclear division^9^. At the center of the septum lies a septal pore that allows the exchange of cytoplasmic constituents between flanking cells^10,11^. In species of the subphylum Pezizomycotina of Ascomycota, these pores can be plugged by the peroxisome-derived, fungal-specific Woronin body^12^. In contrast, species of the subphylum Agaricomycotina of Basidiomycota have developed an endoplasmic reticulum–derived septal pore cap (SPC) to block the pores^13,14^. A mutant lacking the Woronin body matrix protein Hex1 exhibited extensive loss of cytoplasm from flanking cells via septal pores upon hyphal wounding^12^.

Septal pore function is dynamically regulated in response to mechanical, environmental, and physiological conditions. Stresses such as low temperature and low pH reduce cell-to-cell connectivity by unknown mechanisms^10,11^. Hyphal wounding and abiotic stresses induce the accumulation of septal pore-associated (SPA) proteins and SO (or SOFT) protein at the pore^15–17^. Furthermore, the pore closes transiently during mitosis and opens during interphase when Nima kinase, a cell-cycle regulator, moves from the nucleus to the septal pore^18^. Septal pores are also modulated by physiological conditions such as hyphal age^11,19^, suggesting the involvement of additional components in regulating their functions. Considering these dynamic behaviors of septal pore regulation, our understanding of the mechanisms is lacking, particularly given limited knowledge of Woronin body function.

Hyphae, the signature of fungal multicellularity, are proposed to have evolved early in fungal evolution, possibly in the early diverging fungi such as Blastocladiomycota, Chytridiomycota, Zoopagomycota, and Mucoromycota^20^. Using large-scale genomic comparison, Kiss *et al*. proposed a set of genes involved in hyphal morphogenesis that are shared among fungal classes regardless of the presence or absence of filamentous hyphae^20^. Moreover, while fungal species within Pezizomycotina and Agaricomycotina typically show a substantial gain of gene families, as exemplified by morphological complexities such as septal pore, yeasts show a higher rate of gene losses^20,21^. Hyphal morphology is generally considered to be the foundation of fungal multicellularity, while the emergence and regulation of the septal pore have evolved independently in multicellular lineages, conferring further morphological complexities. However, these evolutionary steps of septal pore emergence and regulation remain poorly understood.

Here, we combined genomic comparisons between septal pore–bearing and –lacking ascomycetes with a large-scale localization screening of candidate septal pore proteins to identify novel septal components. We found that the septal pore is a focus for localizing the proteins involved in its dynamic regulation. With the bioinformatics approach including extensive phylogenetic analyses, we elucidated the patterns of gene evolution involved in septal pore plugging upon hyphal wounding.

## RESULTS

### Selection of candidate septal pore proteins

Here, *Aspergillus oryzae* was used for the availability of advanced experimental techniques for quantification of septal pore plugging upon hyphal wounding induced by hypotonic shock, as well as quantifying cell-to-cell connectivity upon cold stress using photoconvertible fluorescent proteins^10,16,19,22,23^. The *A. oryzae* genome was compared with those of septal pore–bearing ascomycetes *A. nidulans*, *A. fumigatus,* and *Neurospora crassa* (Pezizomycotina) as well as septal pore–lacking ascomycetes *Saccharomyces cerevisiae*, *Candida albicans* (Saccharomycotina), and *Schizosaccharomyces pombe* (Taphrinomycotina) using BLASTp (Fig. 1a). Firstly, the proteins known to function in cellular morphogenesis (encoded by 243 genes that have orthologs in *A. fumigatus*^22^) (Supplementary Data 1a) were compared. The majority of the proteins from the septal pore–bearing species showed higher sequence homologies (Fig. 1a, upper blue bubble in bubble chart), and a considerable number of proteins also exhibited sequence homologies with the proteins from the septal pore–lacking species (Fig. 1a, upper bubble chart). This suggested that cellular morphogenesis-related genes evolved before their divergence. Similarly, we analyzed proteins known to function in septal pore regulation (Supplementary Data 1b based on previous reports^15–17,23,24^). The majority of the proteins showed no or limited homologies with yeast proteins (Fig. 1a, lower bubble chart in green), while many proteins exhibited sequence homology in the septal pore–bearing species (Fig. 1a, lower bubble chart in blue). The result suggested that these proteins arose in the lineage leading to Pezizomycotina, but were lost or diverged in ascomycete yeasts. Next, to analyze the conservation of protein–coding genes more widely, we similarly compared whole proteomic datasets from the aforementioned species with that of *A. oryzae*. A large number of proteins with higher sequence homologies were found within the septal pore-bearing species, but a lesser number in the septal pore-lacking species (Fig. 1a, bar graph in blue. This conservation tendency is similar to that of comparative analysis using the proteins involved in septal pore regulation (Fig. 1a, lower bubble chart). Based on these findings, we hypothesized that proteins involved in septal pore regulation might be conserved in septal pore–bearing species, but absent or divergent in the septal pore–lacking species.

**Figure 1.**
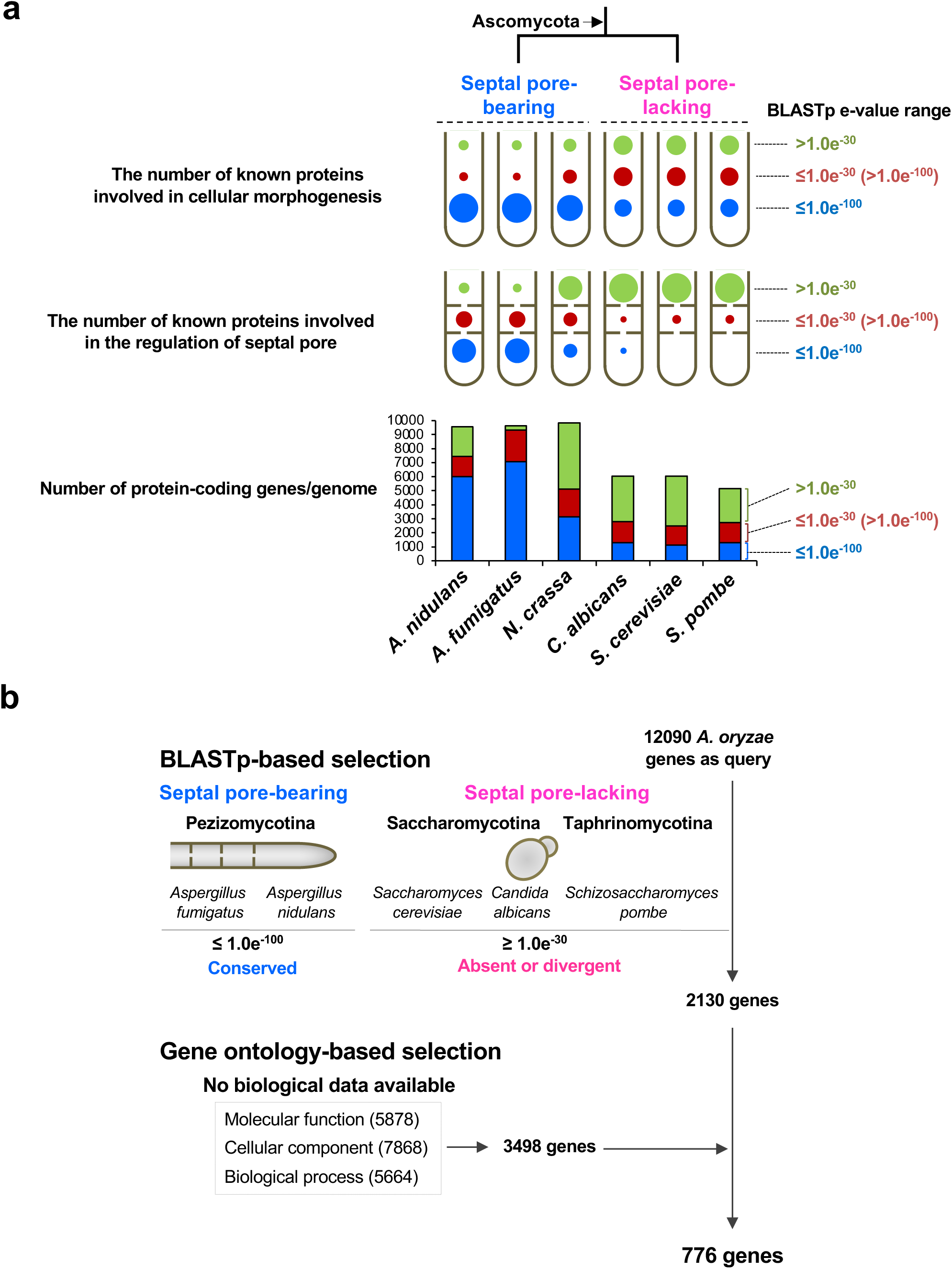
Strategy for selecting candidate septal pore proteins. **(a)** BLASTp-based genomic comparison among the septal pore–bearing and –lacking ascomycetes. Upper cartoons of hyphae show hyphal morphology, and lower cartoons of hyphae with perforated septa indicate septal pore and its regulation. Bubble size is proportional to the number of known proteins involved in hyphal morphology (upper bubble chart, Supplementary Data 1a) and septal pore regulation (lower bubble chart, Supplementary Data 1b). In bar graph, the Y-axis represents the number of total protein-coding genes per analyzed genome. Colors in bubbles and bars represent the proteins within the analyzed BLASTp e-values; blue, red, and green indicate BLASTp e-value ranges ≤ 1.0e-100, ≤ 1.0e-30 (>1.0e-100), and >1.0e-30, respectively. **(b)** Strategy for the selection of candidate septal pore proteins.

To select candidates for septal pore-related proteins, we screened for genes conserved in the septal pore–bearing ascomycetes *A. oryzae*, *A. fumigatus,* and *A. nidulans* (e-values ≤ 1.0e-100 in BLASTp), but absent or divergent in the septal pore–lacking ascomycetes *S. cerevisiae*, *C. albicans*, and *S. pombe* (e-values ≥ 1.0e-30 in BLASTp) (Fig. 1b). A total of 2130 genes were identified as the potential candidates. The resultant 2130 genes were further used for selecting candidates that do not have biological data for any gene ontology (GO) terms; molecular function, cellular component, and biological process (Fig. 1b). Consequently, a subset of 776 uncharacterized candidate genes for septal pore–related proteins (Supplementary Data 2) were finally selected.

### Subcellular localization of candidate proteins

Candidate gene sequences were retrieved from the Aspergillus genome database AspGD (http://www.aspgd.org). As reported in previous global analyses of protein localization in budding^25^ and fission^26^ yeasts, each of the 776 candidate genes was fused to *egfp* at its 3′ end under the control of the inducible *amyB* promoter^27^ (Fig. 2a). The expression cassettes were ectopically inserted into the *niaD* locus of *A. oryzae* strain NSlD1^10^ by homologous recombination (Fig. 2a and Supplementary Data 3: Strain list).

**Figure 2.**
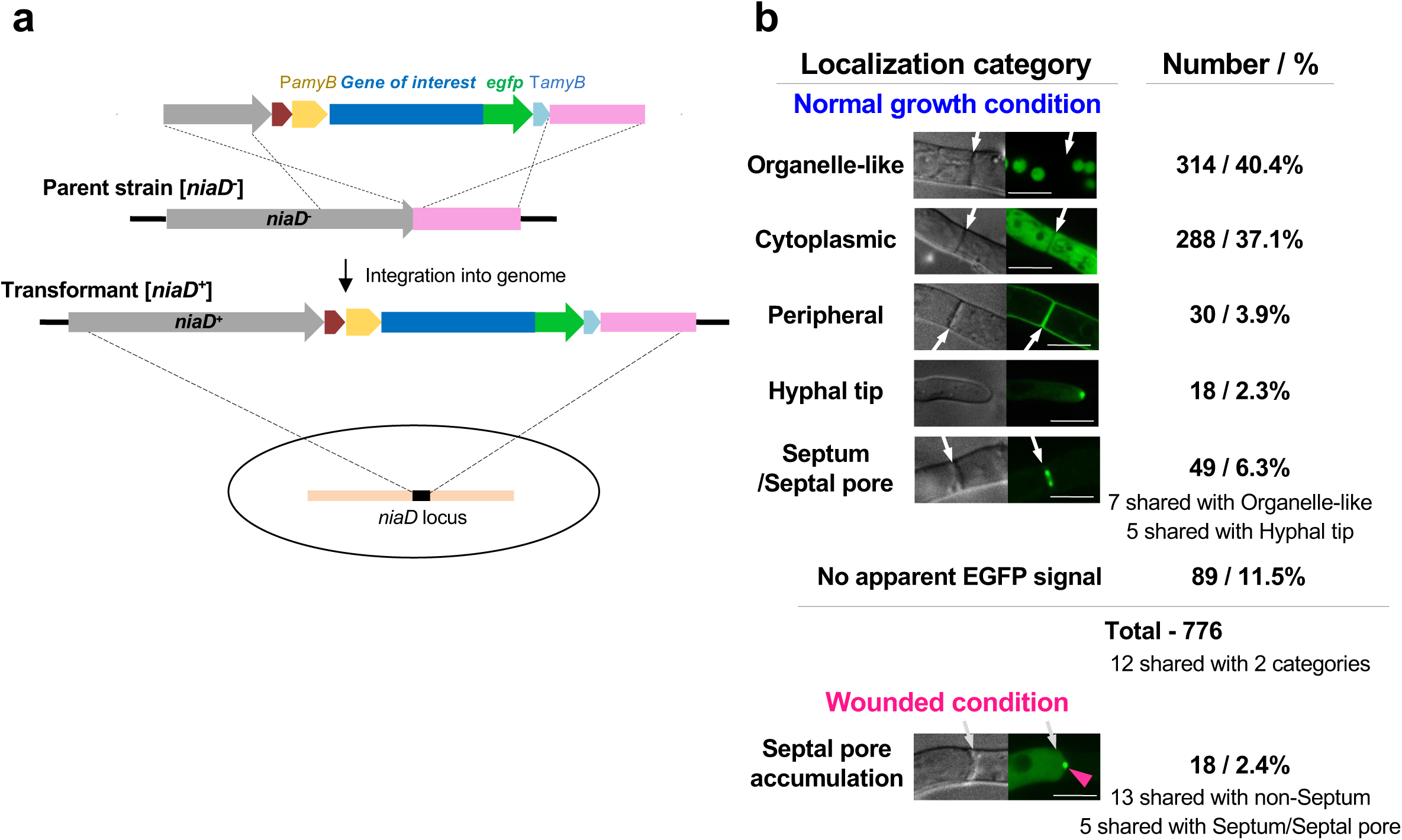
Subcellular localization of candidate septal pore proteins. **(a)** Schematic of the expression cassette for EGFP fusions of selected proteins to be inserted in the *A. oryzae* genome. **(b)** Subcellular localizations of the 776 proteins. Representative confocal fluorescence micrographs of localization categories are shown; arrows indicate septa. Scale bar, 5 μm.

Of the 776 resulting strains, we detected EGFP fluorescence in the live hyphae of 687 (88.5%) strains (Fig. 2b). Their localization patterns were classified into five categories: organelle-like, cytoplasmic, peripheral, hyphal tip, and septum/septal pore (Fig. 2b). A total of 314 (40.5%) proteins predominantly localizing as spherical structures, networks, and puncta resembling cell organelles were categorized as “organelle-like” localization (Fig. 2b and Supplementary Fig. 1). Diffuse, uniform cytoplasmic staining was observed in 288 (37.1%) proteins, and 30 (3.9%) exhibited peripheral localization (Fig. 2b and Supplementary Figs. 2, 3a). However, many of the peripheral proteins overlapped with organelle-like distributions (Supplementary Fig. 3a), probably due to their presence in the secretory pathway. Nineteen of the peripherally localized proteins contained signal peptides and/or transmembrane domains (Supplementary Fig. 3a), supporting this hypothesis. Eighteen (2.3%) proteins localized to the hyphal tip (Fig. 2b and Supplementary Fig. 3b), whereas the remaining 89 proteins (listed in Supplementary Data 4) did not yield detectable EGFP signals.

**Figure 3.**
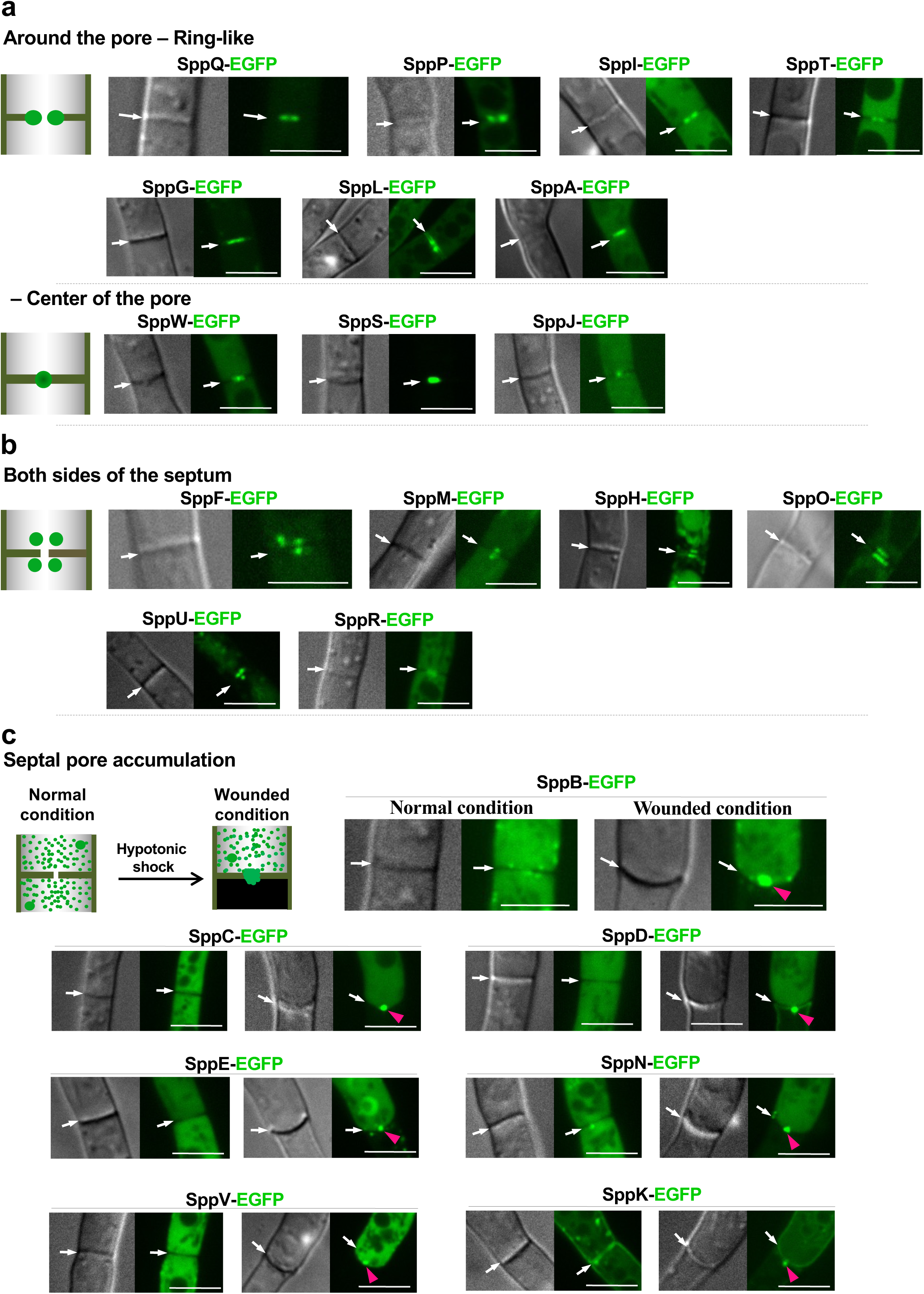
Localization of SPP proteins to the septum and septal pore. The apical septa of strains expressing EGFP–fused proteins were analyzed by confocal fluorescence microscopy under normal, control growth conditions and upon hyphal wounding. Cartoons represent the localization pattern categories observed at the septum. **(a)** Localization around the septal pore. **(b)** Localization on both sides of the septum. **c** Accumulation of non-septal SPP proteins at the septal pore upon hyphal wounding induced by hypotonic shock. White arrows indicate septa and red arrowheads indicate septal-pore accumulation. Scale bars, 5 μm.

Importantly, 62 proteins (8.0%) localized to the septum or septal pore (Fig. 2b). Forty-nine of them were septal under normal growth conditions (Fig. 3ab and Supplementary Fig. 4a-c), and the remaining 13 accumulated at the septal pore upon hyphal wounding (Fig. 3c and Supplementary Fig. 4d). Interestingly, five septal proteins had overlapping distributions with those of hyphal tip category (Supplementary Fig. 3b), suggesting regulatory networks shared between the septum and hyphal tip.

Collectively, our bioinformatics-based screening strategy allowed to find many proteins showing the localization related to the septum.

### Septal pore is the subcellular site to which many proteins localize

Of the 62 septal proteins, we later found that 23 were functionally involved in septal pore plugging upon hyphal wounding (see Fig. 4b) and designated them as <u>s</u>eptal <u>p</u>ore <u>p</u>lugging (SPP) proteins with alphabetical order of functional importance in the septal pore-plugging activity (Fig. 4b).

**Figure 4.**
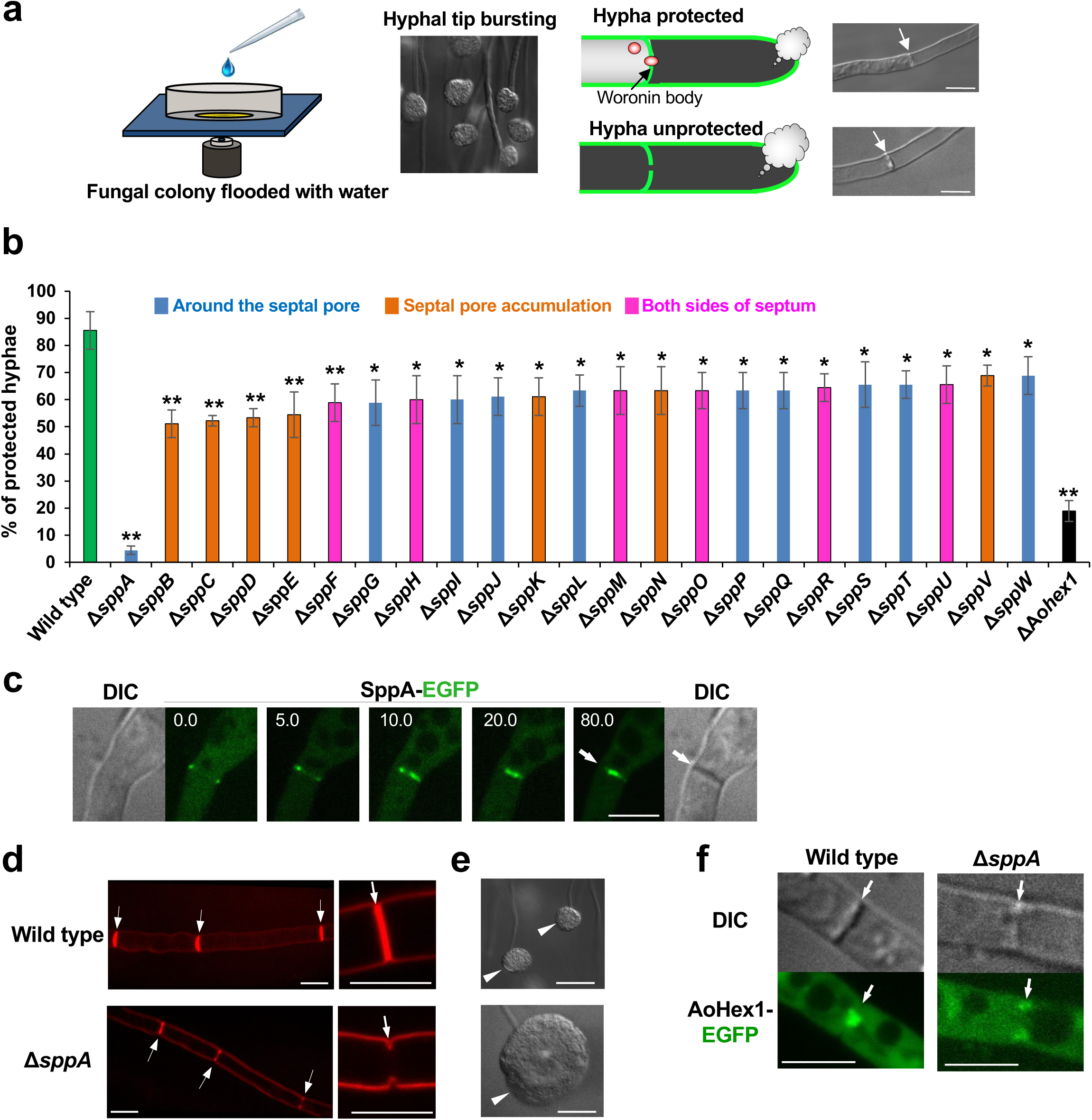
Regulation of septal pore plugging by SPP proteins upon wounding. **(a)** Schematic showing hyphal tip bursting induced by hypotonic shock. Cartoons and DIC images show wounded hyphae capable (top) or incapable (bottom) of protecting adjacent cells from cytoplasmic loss. Scale bars, 5 μm. **b** Protection of flanking cells from excessive loss of cytoplasm upon hyphal wounding. Thirty randomly selected hyphae showing hyphal tip bursting were observed in each experiment. Three independent experiments were performed, and the percentage of hyphae protected from the excessive loss of cytoplasm is shown in the Y axis. Error bars represent standard deviations. *\*p <* 0.05; *\*\*p <* 0.01 (Student’s *t-*test). **c** SppA localization at the site of septum formation. Strains were grown for 8 h in CD (2% glucose) liquid medium supplemented with 1% casamino acids. Time is indicated in minutes. White arrows indicate septa. Scale bars, 5 μm. **d** Incomplete septum formation in Δ*sppA* visualized with FM4-64. White arrows indicate septa. Scale bars, 5 μm. **e** Leaked cytoplasmic constituents in Δ*sppA* upon hyphal tip bursting induced by hypotonic shock. White arrowheads indicate leaked cytoplasmic constituents at the hyphal tip. Scale bars, 20 μm. **f** Abnormal tethering of Woronin bodies to the incomplete septum in Δ*sppA*. Single confocal fluorescence images at apical septa are shown. White arrows indicate septa. Scale bars, 5 μm.

Under normal growth, 49 proteins were classified into three categories based on their septal localizations: 1) around the septal pore (Fig. 3a and Supplementary Fig. 4a), 2) on both sides of the septum (Fig. 3b and Supplementary Fig. 4b), and 3) along the septum (Supplementary Fig. 4c).

The first category was further divided into two subgroups: ring-like localization, as two dots around the septal pore under confocal microscopy (26 proteins); and focal localization at the center of the septal pore (eight proteins) (Fig. 3a and Supplementary Fig. 4a).

Thirteen proteins were present on both sides of the septum near the septal pore, similar to the location of the Woronin body (Fig. 3b and Supplementary Fig. 4b). SppF and SppM showed punctate localization on both sides, parallel to the septal pore. While the rest of the proteins under this category localized broadly to both sides with multiple puncta (Fig. 3b and Supplementary Fig. 4b). SppF-EGFP colocalized with mCherry–tagged Woronin body matrix protein AoHex1 at the septum and in the cytoplasm (Supplementary Fig. 5a). However, the location of the Woronin body to the septum was independent of SppF (Supplementary Fig. 5b). Two proteins were localized in the third category, along the septum but not peripherally along the hyphae (Supplementary Fig. 4c).

The interconnected array of hyphal cells is vulnerable to mechanical injury, causing extensive loss of cytoplasm in the absence of septal pore plugging^12,14,18,24,25^. A number of cytoplasmic proteins accumulate at the septal pore upon wounding^12,17,18^. Therefore, all 776 EGFP fusion-expressing strains were subjected to hypotonic shock–induced hyphal wounding^12,18,24,25^. Interestingly, 13 proteins showing cytoplasmic (five proteins), organelle-like (four proteins), and hyphal tip (four proteins) localizations under normal growth conditions were found to accumulate at the septal pore upon wounding (Fig. 3c and six proteins in Supplementary Fig. 4d). Furthermore, SppI, SppT, and SppL, which localized around the septal pore, as well as SppO and SppF, which localized on both sides of the septum under normal growth, showed concentrated localization to the septal pore upon wounding (Supplementary Fig. 4e). The results highlighted them as candidate proteins with functions related to the response to wounding, as reported for the septal pore–accumulating proteins^17–19^.

### Role of SPP proteins in septal pore plugging

In animals, mechanical wounding induces dynamic remodeling of membrane function via mechanisms including fusion of secretory vesicles to the plasma membrane to promote healing^28^. Pezizomycotina employs the Woronin body to plug the septal pore upon wounding^12^. Here, septal pore-plugging activity was analyzed after the deletion of 62 candidate genes by replacing them with the *pyrG* selectable marker. Four deletions exhibited reduced colony growth (Supplementary Fig. 6a). The deletion strains were subjected to hypotonic shock, which causes hyphal tip wounding (Fig. 4a)^10,16,22,23^. Hyphae protected from excessive cytoplasmic loss from the adjacent cells were counted using differential interference contrast microscopy (Fig. 4a).

In total, 23 (37%) deletion strains showed a significantly decreased ability to protect flanking cells from excessive cytoplasmic loss upon wounding compared with wild-type control (Fig. 4b), indicating impaired septal pore plugging. Notably, deletion of *sppA* did not prevent cytoplasmic loss (Fig. 4b). This inability is possibly due to a defect in septum formation, as SppA-EGFP transiently appeared at the site of septum formation, similar to contractile actin ring assembly and constriction (Fig. 4c). SppA, which possesses the C2 domain (Fig. 6), is the ortholog of Inn1 and Fic1 in budding and fission yeasts, respectively (Supplementary Fig. 7a), both of which are essential for plasma membrane ingression during cytokinesis^29,30^. Because septum formation in Pezizomycotina is analogous to cytokinesis, we evaluated septum morphology in Δ*sppA*. Wild-type controls exhibited disc-shaped walls compartmentalizing the hyphae (Fig. 4d). In contrast, Δ*sppA* showed abnormal septa with incomplete ingression of the plasma membrane from the cortex (Fig. 4d). A larger volume of cytoplasmic constituents from continuously interconnected cells leaked in Δ*sppA* upon hyphal wounding (Fig. 4e). In addition, Woronin bodies were abnormally tethered to the incomplete septum at the hyphal cortex in Δ*sppA* (Fig. 4f).

The absence of SppB, SppC, SppD, SppE and SppF increased cytoplasmic loss via the septal pore relative to that in wild-type strain (Fig. 4b), indicating the functional relevance of these proteins in septal pore plugging upon wounding. Expression of EGFP-fusion genes attenuated the septal pore-plugging defect of the corresponding gene deletions (Supplementary Fig. 6b), confirming that the fusions are functional. We visualized Woronin bodies in the gene deletion backgrounds and found that they remained tethered to the septum (Supplementary Fig. 5b), suggesting that their tethering occurs independently of the SPP proteins. However, over half of the 62 analyzed proteins did not appear to play a significant role in septal pore plugging upon wounding.

To investigate functional redundancy among SPP proteins, we generated double deletion strains. Representative genes *sppG*, *sppF*, and *sppE* were selected from localization categories; around the septal pore, on both sides of septum, and septal pore accumulation, respectively. The selected representative genes were deleted in the deletion background of another gene under the same localization category. After hyphal wounding, none of the double deletions except for Δ*sppG*Δ*sppA* showed a further reduction in septal pore plugging compared with that of the single deletion (Supplementary Fig. 6c). Moreover, the single deletions revealed a measurable phenotype, whereas the corresponding double deletions did not display further deficiencies in septal pore plugging. These results suggested that each SPP protein played a non-overlapping role and could not be fully substituted by one another.

### SPP proteins and cell-to-cell connectivity under cold stress

Because the septal pore closes under stresses^10,11^, 62 strains expressing the septum-localizing proteins tagged with EGFP were subjected to cold stress (4°C) to assess the dynamic behavior at the septal pore. Interestingly, four SPP proteins that normally localized to cytoplasm were observed at the septal pore under cold stress (Fig. 5a). SppB, SppE, and SppN accumulated at the septal pore from both flanking cells, while SppC was present as puncta at the center of the septal pore (Fig. 5a). Localization of SppJ to the septal pore was promoted by cold stress (Fig. 5a).

**Figure 5.**
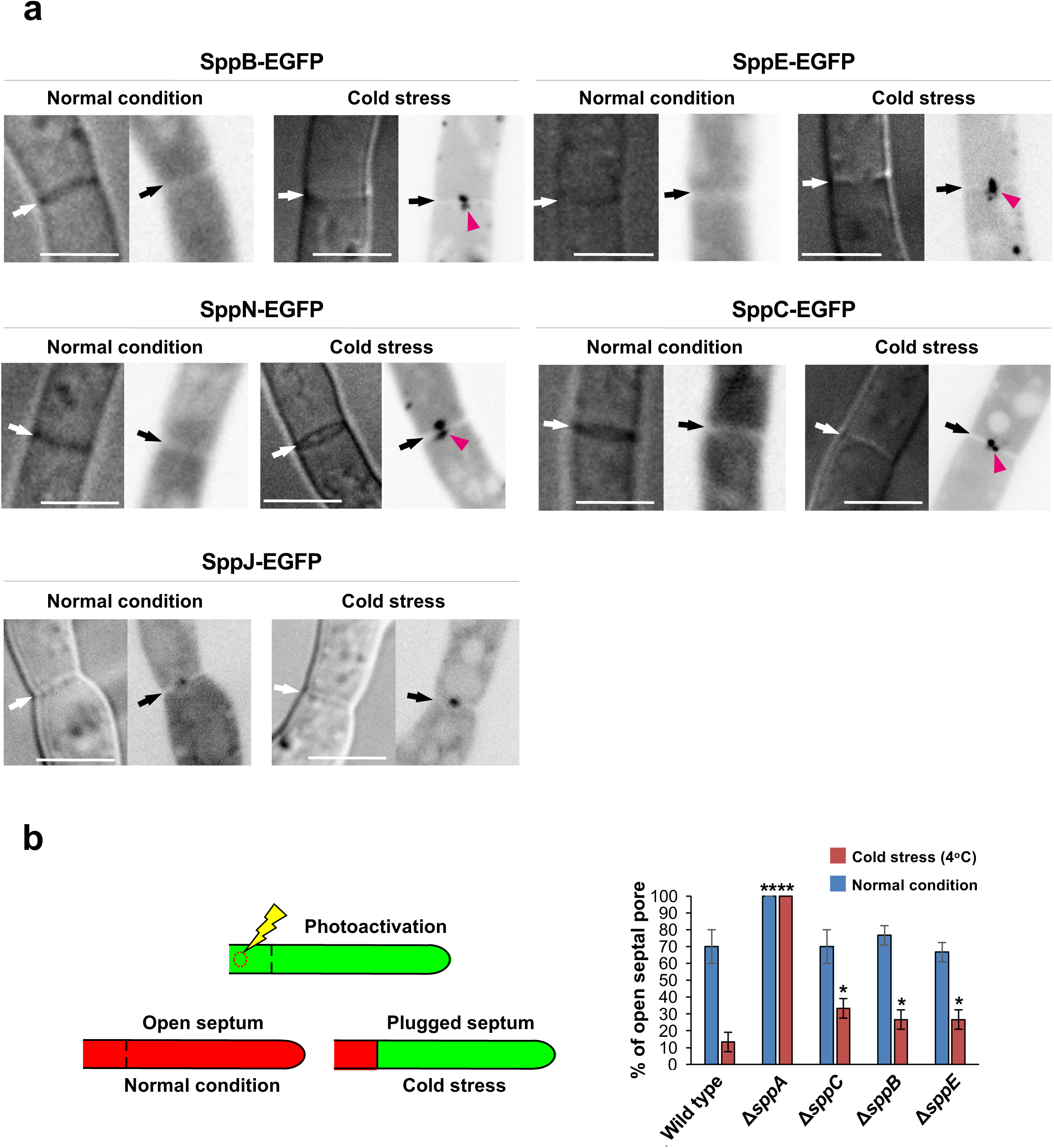
Regulation of cell-to-cell connectivity by SPP proteins under cold stress. **a** Localization of SPP-EGFP fusion proteins. Black/white arrows indicate septa and pink arrowheads indicate septal pore accumulation. Single confocal fluorescence images are shown. Scale bars, 5 μm. **b** Models for photoactivation and cell-to-cell transfer of Dendra2 (left); quantification of transfer (right). Dendra2 transfer through the septal pore was monitored at 10 randomly chosen apical septa in each experiment. Bars, 10 μm. *n* = 3. Error bars = *SD*. *\*p* < 0.05 *\*\*p* < 0.01 (Student’s *t-*test).

The status of cell-to-cell connectivity in the gene deletion backgrounds lacking 62 septal proteins was monitored under cold stress by tracking the cell-to-cell transfer of Dendra2, a green-to-red photoconvertible fluorescent protein^10^ (Fig. 5b left). In wild-type controls, the red fluorescence of photoconverted Dendra2 was transferred to adjacent cells via the septal pore, but this transfer was dramatically decreased at 4°C (Fig. 5b, right). In contrast, the diffusion of Dendra2 was not restricted at all in Δ*sppA* under both normal and cold temperatures (Fig. 5b, right), indicating continuous cell-to-cell connectivity caused by incomplete septum formation (Fig. 4d). The Δ*sppB*, Δ*sppE,* and Δ*sppC* strains still allowed Dentra2 transfer at low temperature (Fig. 5b right), indicating aberrant regulation of the septal pore function upon cold stress.

### Role of the disordered region of SPP proteins in septal pore accumulation

A large number of septal proteins including SPP proteins are associated with a tiny subcellular site, the septal pore. Next, we elucidated whether the SPP proteins possess specialized sequences or structural features, which mechanically target them to the septum as reported with the disordered region of Spa proteins^19^. Firstly, we analyzed the degree and distribution of disordered regions present in the SPP proteins using IUPred2A^31^. Eleven of the 23 SPP proteins possess sequences with a high probability of intrinsic disorder (Fig. 6 and Supplementary Fig. 7a-b). Typically, SPP proteins localizing on both sides of the septum were structurally ordered, while those localizing around or accumulating at the septal pore were mostly disordered (Fig. 6 and Supplementary Fig. 7b). To investigate the importance of the disordered region in septal localization, we expressed EGFP-tagged truncated variants. Interestingly, the disordered N-termini of both SppB and SppD were sufficient and essential for septal accumulation upon wounding (Fig. 6a). Disordered regions containing proline-rich domains are important for protein aggregation in phase separation behavior^32,33^. SppN sequence was prone to having disordered region across its entire length and contained conserved proline-rich motifs in the N-terminal residues 1-265 (Supplementary Fig. 7c-d) that were sufficient and required for wound-induced septal accumulation (Fig. 6a). Contrastingly, in SppP, SppI, SppL, and SppW, both the ordered and disordered regions were essential for their normal septal localization (Fig. 6b). Collectively, the disordered region is important for the wound-induced septal accumulation and normal septal localization. Next, we analyzed the disordered regions in terms of amino acid composition and compared them with those of *N. crassa* SPA proteins, phenylalanine/glycine (FG)-repeat nucleoporins (FG-Nups), serine/arginine (SR)-repeat splicing factors, and disordered proteins from the DisProt dataset. Both SPP and SPA proteins revealed biases to arginine and histidine but showed antipathies to lysine and glycine when compared with FG, SR, and Disprot dataset (Supplementary Fig. 7c).

**Figure 6.**
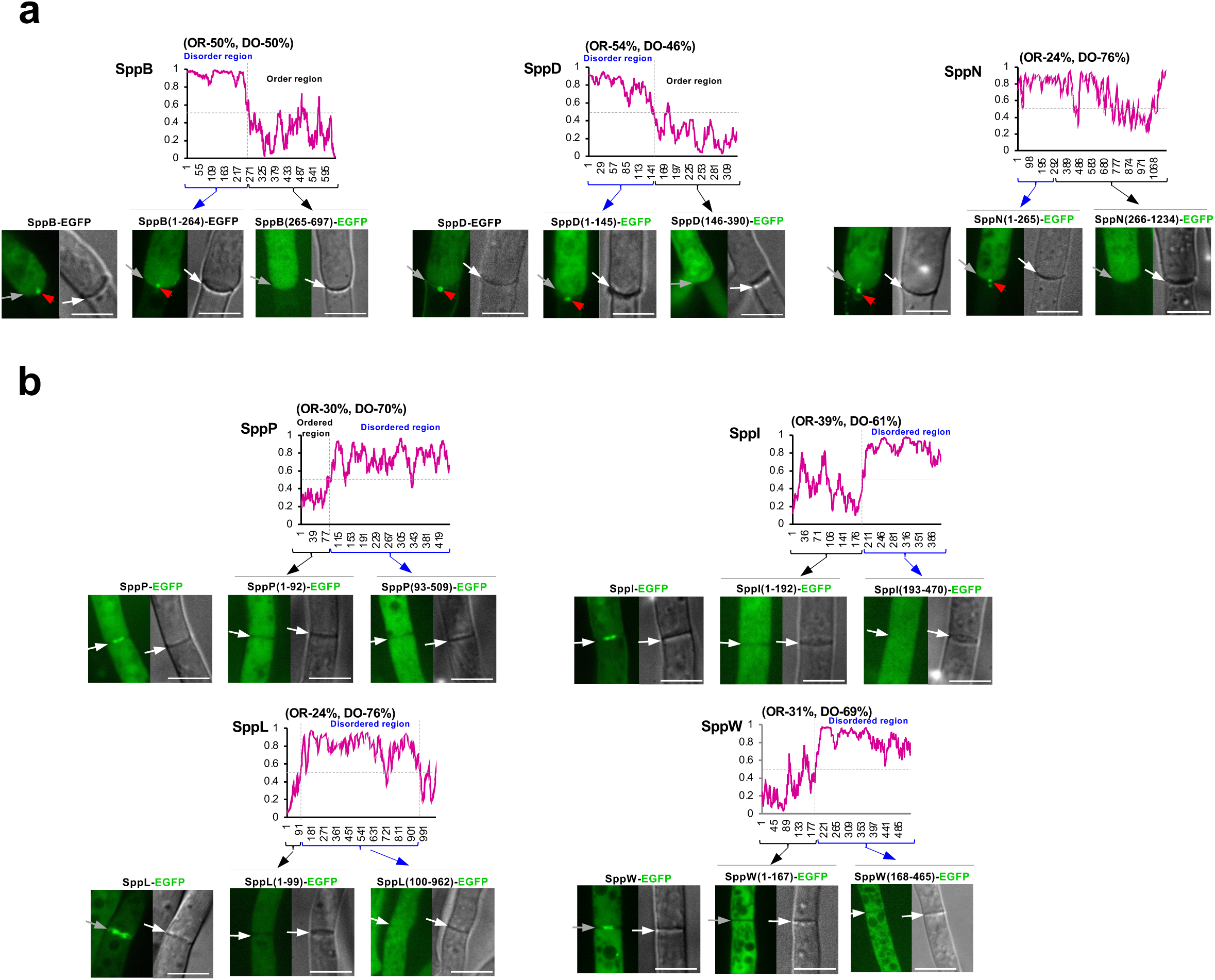
Prediction of disordered regions and analysis of truncated SPP protein variant localization. **a** Three SPP proteins showing septal pore accumulation upon hyphal wounding harbored a considerable length of a disordered region separated from the ordered region. **b** Four SPP proteins localized around the septal pore and harbored a considerable length of the disordered region separated from the ordered region. Graphs show the prediction of disordered regions using IUPred2A; the Y-axis indicates the predicted probability of disorder and the X-axis represents the amino acid sequence. In the fluorescence microscopic analysis, full-length and truncated variants of EGFP–fused SPP proteins were expressed in the corresponding deletion background. Strains were grown in CD medium supplemented with 1% casamino acids for 18 h. Single confocal fluorescence images are shown. Arrows indicate apical septa, and red arrowheads indicate septal pore accumulation. Scale bars, 5 μm. OR, ordered; DO, disordered.

### Orthologous relation of SPP proteins in fungi

We analyzed the phylogenetic distribution and the degree of divergence of SPP proteins compared with broad data of protein sequences retrieved from representative species covering all major fungal phyla and subphyla. Accordingly, we selected species from the subphyla within Ascomycota (Taphrinomycotina, Saccharomycotina, and Pezizomycotina) and Basidiomycota (Agaricomycotina, Pucciniomycotina, and Ustilaginomycotina), as well as representatives of early diverging Mucoromycota, Zoopagomycota, Chytridiomycota, Blastocladiomycota, and Cryptomycota (Rozellomycota)^20,34^. Due to its high rate of evolution^34^, Microsporidia was excluded from this analysis. First, proteome data of 81 fungal species were categorized into orthologous groups using OrthoFinder^35^. Subsequently, substitution rates of orthologous proteins in fungal phyla/subphyla from the Pezizomycotina root were calculated based on maximum likelihood phylogenetic trees (Fig. 7 and Supplementary Data 5a-b). To further verify orthologous relations, SPP protein phylogenies were generated for each orthologous group (Supplementary Figs. S8 and 9; white dots in Fig. 7). Together with phylogenetic analyses, SPP proteins were finally classified into two evolutionary groups; 1) those with orthologs outside of Pezizomycotina, and 2) those present specifically in Pezizomycotina.

**Figure 7.**
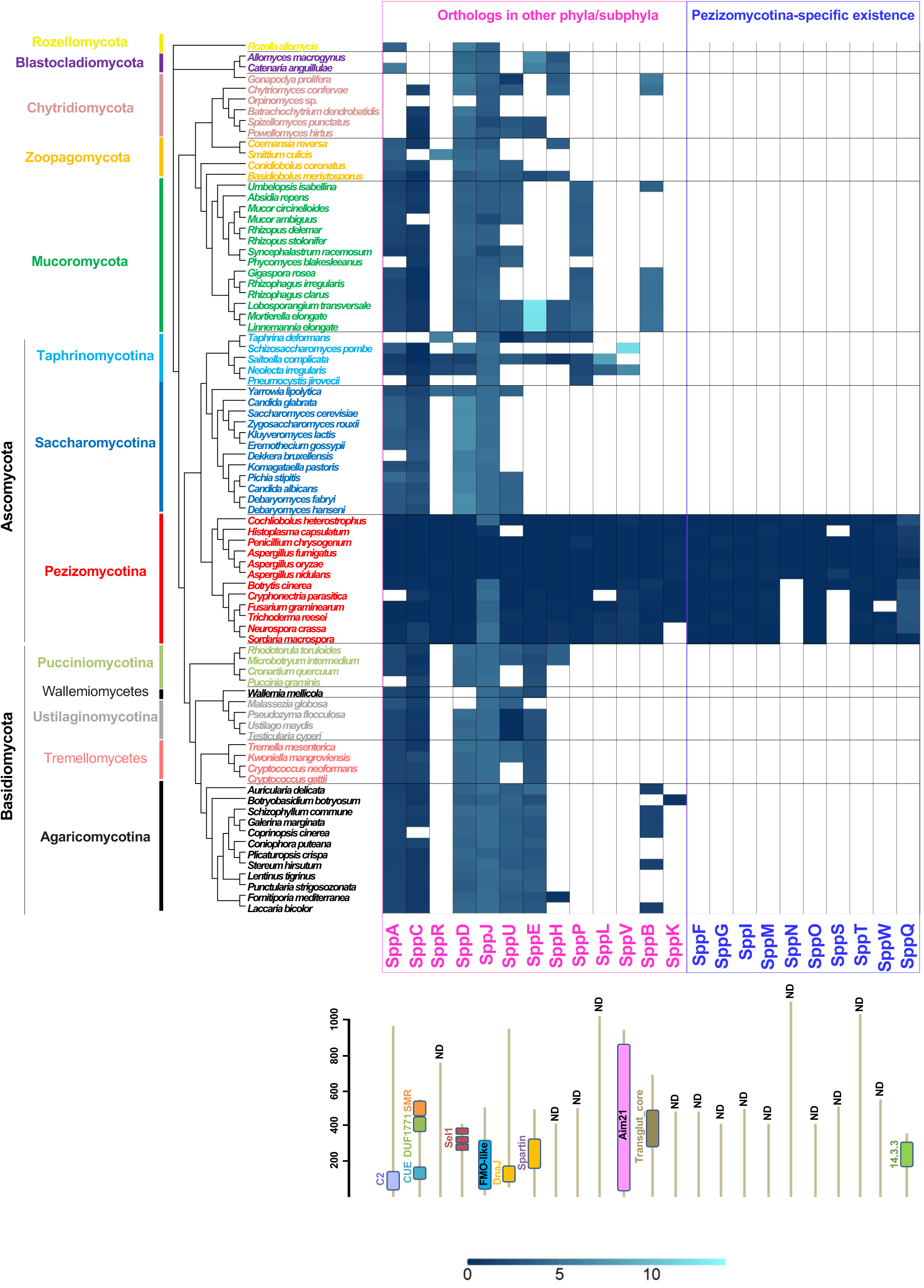
Orthologous relations and substitution rates of SPP proteins among fungal taxa. Amino-acid substitutions per site are shown in blue as a heat map. Filled and empty colors denote the presence and absence of orthologous proteins, respectively, according to the results of OrthoFinder. The phylogenetic tree was generated using the orthologous group proteins with a single ortholog present in almost all the species. The scale at the left of the models of SPP protein structures represents numbers of amino acids. ND, no domain.

Thirteen proteins (SppA, SppC, SppR, SppD, SppJ, SppU, SppE, SppH, SppP, SppL, SppV, SppB, and SppK) exhibited orthologs outside of Pezizomycotina. Among them, SppA, SppC, and SppR possessed orthologs in septal pore–lacking ascomycete yeasts Saccharomycotina and Taphrinomycotina in the same clade. SppA contains the C2 domain (PF00168)^36^ and belongs to a broad orthologous group that includes Inn1 and Fic1 from budding and fission yeasts, respectively (Fig. 7 and Supplementary Fig. 8a). SppC possesses an N-terminal CUE (PF02845)^37^, a C-terminal SMR (PF01713)^38^, and DUF1771 (SM001162)^39^ domains. Similar to SppA, SppC belongs to a broad orthologous group (Fig. 7 and Supplementary Fig. 8b). SppR was grouped with the peroxisomal membrane protein Pex8 from some fungal species (Fig. 7 and Supplementary Fig. 8c).

SppD, SppJ, and SppU exhibited multiple subclades, including Pezizomycotina, within their orthologous groups. SppD possesses three repeats of the SEL1 domain (SM00671)^40^ and forms a Pezizomycotina-specific subclade within the large clade including other fungal orthologous proteins (Fig. 7 and Supplementary Fig. 8d). SppJ contains a DnaJ domain (PF00226)^41^ and forms a Pezizomycotina-specific subclade close to some of the species from Chytridiomycota and Mucoromycota (Fig. 7 and Supplementary Fig. 8e). SppU contains an FMO-like domain (PF00743)^42^ and forms a subclade including Pezizomycotina and a limited number of Ustilaginomycotina, Taphrinomycotina, and Chytridiomycota species (Fig. 7 and Supplementary Fig. 8f).

Furthermore, SppE, SppH, SppP, SppL, and SppV proteins exhibited orthologous relations with other fungal phyla/subphyla, including the ascomycete yeasts Taphrinomycotina. SppE contains a plant-related senescence domain (PF06911)^43^ and belongs to an orthologous group including Basidiomycota, early diverging fungi, and Taphrinomycotina (Fig. 7 and Supplementary Fig. 8g). SppH and SppP, which do not possess known domains, shared orthologous relations with proteins from early diverging fungi and some species of Taphrinomycotina (Fig. 7 and Supplementary Fig. 8h-i). SppL and SppV, with latter harboring an Aim21 domain (PF11489)^45^, exhibited orthologous relations with a limited number of species from Taphrinomycotina (Fig. 7 and Supplementary Fig. 8j-k).

SppB and SppK also exhibited orthologs outside of Pezizomycotina, but not in any of the ascomycete yeasts. SppB, which contains a Transglut_core domain (PF01841)^44^, belonged to an orthologous group that includes some species from Chytridiomycota, Mucoromycota, and Agaricomycotina (Fig. 7 and Supplementary Fig. 8l). SppK showed an orthologous relation only with some species from Agaricomycotina (Fig. 7 and Supplementary Fig. 8m).

As the other group, a total of 10 SPP proteins were identified as Pezizomycotina-specific existence. Nine SPP proteins, SppF, SppG, SppI, SppM, SppN, SppO, SppS, SppT, and SppW, do not harbor any domains but lacked orthologous relations with other fungal phyla and subphyla (Fig. 7 and Supplementary Fig. 9a-h). SppQ contains a 14-3-3 domain (PF00244)^46^ and belongs to an orthologous group containing Pezizomycotina species only, which is distinct from another group containing the same domain (Fig. 7 and Supplementary Fig. 9i). These SPP proteins are conserved commonly within Pezizomycotina with exceptions for SppN and SppS (Fig. 7).

### Functional dissection of SppA and SppC

Despite our bioinformatics-based screen to select genes absent or divergent in ascomycete yeasts (Fig. 1b), two SPP proteins SppA and SppC contain respective yeast orthologs within the same clades (Supplementary Fig. 8a, b). Therefore, these proteins were functionally analyzed based on their structural differences.

SppA possesses an N-terminal C2 domain, which is conserved in the fungal orthologous proteins. However, the length of C-terminus varies among the orthologous proteins and is longer in Pezizomycotina species than ascomycete yeasts and other fungi (Fig. 8a). To elucidate the functional importance of individual regions, truncated variants of SppA were expressed as EGFP fusions in a Δ*sppA* background. Their expression was equivalent to that of the full-length SppA (Supplementary Fig. 10a left). An SppA variant (residues 128-912) lacking the N-terminal region along with C2 domain could neither complete septum formation nor prevent excessive cytoplasmic loss upon hyphal wounding (Fig. 8b-d). In addition, this variant did not localize to the septum (Supplementary Fig. 10b), which is consistent with the known function of membrane targeting by the C2 domain^36^. Similarly, SppA truncations lacking the C-terminus (residues 1-127 and 1-476) failed to attenuate the septum pore-defective phenotype caused by s*ppA* deletion (Fig. 8bd), indicating the requirement of these regions for proper septum formation and septal pore plugging. Next, we fused the N-terminal region of SppA along with the C2 domain of *A. oryzae* and the C-terminal region of septal pore–bearing *A. nidulans* and *N. crassa,* as well as of septal pore–lacking *S. pombe* (Fig. 8c). EGFP fusions of these chimeric constructs were expressed similarly to the full-length SppA (Supplementary Fig. 12a left). Chimeric proteins containing the C-termini from *A. nidulans* [SppA(AO+AN)] and *N. crassa* [SppA(AO+NC)] rescued the septal pore-defective phenotypes completely and partially, respectively (Fig. 8c-d). In contrast, the construct with the C-terminus from the *S. pombe* ortholog Fic1 [SppA(AO+SP)] exhibited a defect similar to that of Δ*sppA* (Fig. 8cd), suggesting that the yeast protein is not a completely functional ortholog of SppA. Therefore, we concluded that the extended C-terminal region of SppA provided septum-related function(s) that have diversified the lineage leading to the Pezizomycotina, or earlier but was shortened in ascomycete yeasts.

**Figure 8.**
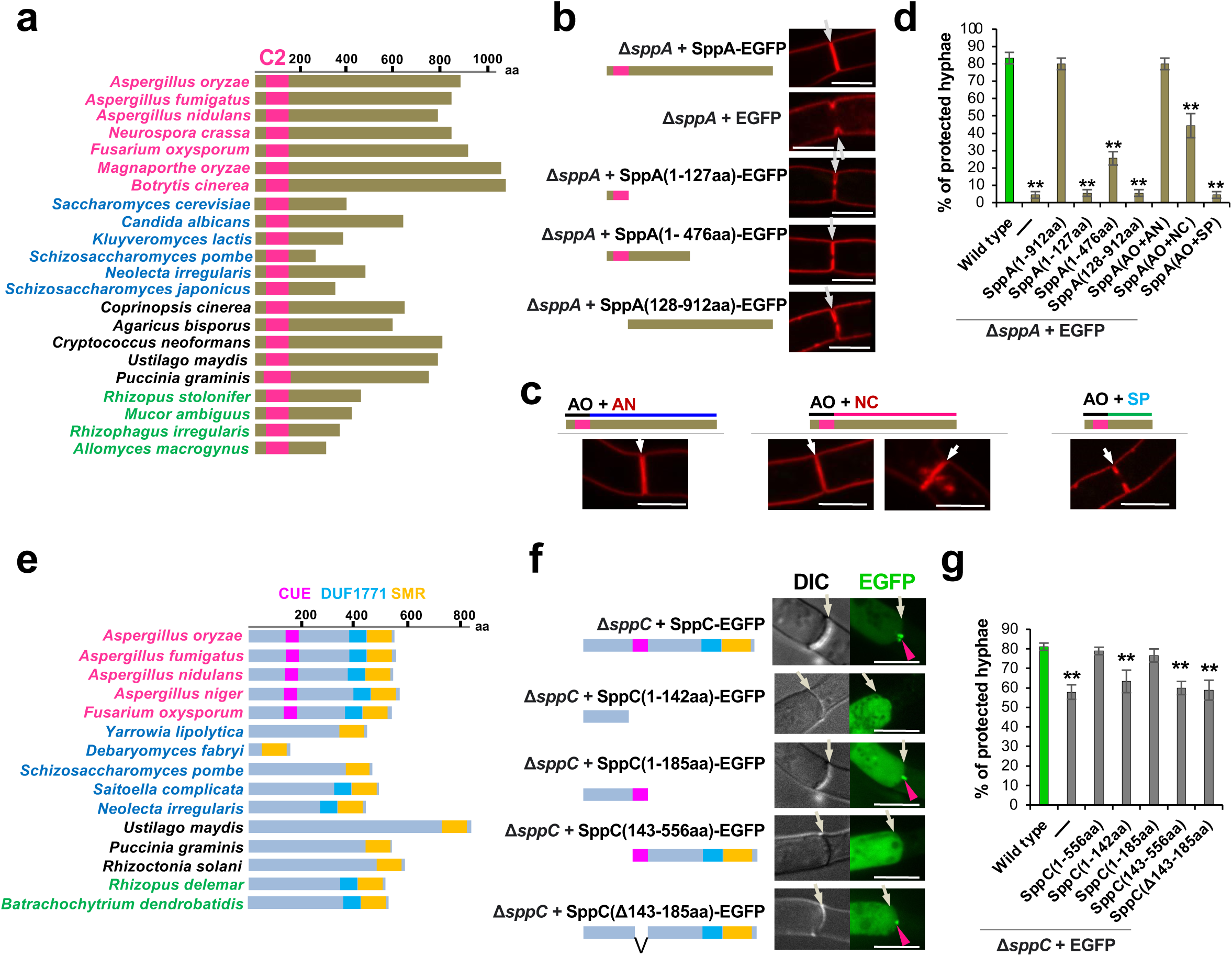
Functional dissection of SppA and SppC. **a** and **e** Lengths of SppA and SppC with their fungal orthologs. Pink, blue, black, and green texts indicate species in Pezizomycotina, ascomycete yeasts, Basidiomycetes, and early diverging fungi, respectively. Bars represent known protein domains. Numbers of amino acid residues are shown in the scale at the top. **b** Both N-and extended C-terminal regions of SppA are essential for normal septum formation. Apical septa were visualized using FM4-64 dye. **c** Septa in strains expressing SppA chimeric proteins with the N-terminus along with the C2 domain derived from *A. oryzae* and the C-terminus derived from the septal pore–bearing or –lacking ascomycetes. AO, *Aspergillus oryzae*; AN, *Aspergillus nidulans*; NC, *Neurospora crassa*; SP, *Schizosaccharomyces pombe*. Septa were visualized using FM4-64 dye. SppA(AO+NC) rescued the defective phenotypes to a notable extent and the two typical images of normal and abnormal septum formation are shown. Scale bars, 5 μm. **d** and **g** Protection of flanking cells from cytoplasmic loss upon hyphal wounding. Error bars represent standard deviations. *\*\*p <* 0.01 (Student’s *t-*test). **f** The N-terminus of SppC is essential for the accumulation at the septal pore upon hyphal wounding. Single confocal fluorescence images are shown. White arrows indicate septa and red arrowheads point to the septal pore localization. Scale bars, 5 μm.

SppC possesses an N-terminal CUE domain, which shows diversity of the domain presence or absence among its orthologs (Fig. 8e and Supplementary Fig. 8b). Multiple sequence alignment revealed diversity in MFP motifs but conservation of the LL motif (Supplementary Fig. 11), both of which are required for the high-affinity binding with ubiquitin^47,48^. To identify the regions responsible for the septal pore-targeted accumulation, truncations of SppC were expressed as EGFP fusions in the Δ*sppC* background. Their expression was equivalent to that of the full-length SppC (Supplementary Fig. 10a right). The N-terminal 185 residues containing the CUE domain were sufficient to accumulate at the septal pore upon wounding, similar to the full-length protein, while truncation of the N-terminal 142 residues prevented this response (Fig. 8f). However, only the N-terminal 142 residues without CUE domain did not clearly accumulate at the septal pore (Fig. 8f), indicating that the N-terminal 142 residues are essential but alone not sufficient to accumulate at the septal pore. A longer N-terminal construct (residues 1-185) containing the CUE domain fully rescued the septal pore-defective phenotype of the Δ*sppC* (Fig. 8g). Two other SppC truncated variants (residues 143-556 and CUE domain–lacking Δ143-185) did not rescue the defective phenotype (Fig. 8g). Collectively, these results indicated that a complete N-terminus including the CUE domain was essential for the septal pore-related function of SppC.

Taken together, these results provide evidence that Pezizomycotina-specific features, such as the extended C-terminus of SppA and the N-terminal region of SppC, are important for septal function.

## DISCUSSION

In this study, we compared two morphologically diverse fungal groups to better understand the genetic reconstruction leading to the evolution of fungal septal pore and its regulation. Localization screening identified numerous functionally important septal proteins that were either absent or divergent in septal pore–lacking yeasts. Our findings show that the Pezizomycotina have evolved at least 23 SPP proteins, which are classified either as those with orthologs outside of Pezizomycotina, or as those present specifically to Pezizomycotina, for septal pore regulation against adverse conditions such as wounding and cold stress.

Though large-scale localization screenings have previously been used as a powerful reverse-genetic approach in animals^49^, yeasts^25,26^, and bacteria^50^, here we for the first time employed the strategy in multicellular fungi. Proteins localizing at the hyphal tip were found less frequently than those localizing at the septum (Fig. 2b and Supplementary Fig. 3b). This could be because some ascomycete yeasts, including the species used for our bioinformatics-based screening (Fig. 1b), can undergo pseudohyphal or hyphal growth^51^. Therefore, Pezizomycotina has not evolved many proteins that function specifically in hyphal tip growth. Almost half of the hyphal tip proteins also localized to the septum/septal pore (Supplementary Fig. 3b), similar to several known septal proteins^10,24^. Of them, three deletion strains lacking such proteins showed reduced colony growth compared to the wild-type strain (Supplementary Fig. 6a; except for SppA), suggesting their involvement in hyphal morphogenesis. The perforation of the septum after its formation/cytokinesis is evolutionarily specific to and emerged uniquely in multicellular fungi. Over half of our identified septal proteins (34/62) exhibited localization around the septal pore (Fig. 3a and Supplementary Fig. 4a), possibly contributing to septal-pore structure maintenance.

Filamentous fungi have evolved wound-management systems with multicellularity. The septate subphyla Pezizomycotina and Agaricomycotina have the Woronin body and the septal pore cap, respectively, to limit cytoplasmic leakage upon wounding^12–14^. In contrast, the aseptate phylum Mucoromycota exhibits protoplasmic gelation^52^. Here, quantitative analysis of protected hyphae upon wounding revealed that more than a third (23/62) of the candidate septal proteins were involved in septal pore plugging (Fig. 4b). Characterization of the functional domains present in several SPP proteins will help to address conservation of regulatory components/mechanisms across the cell-to-cell connectivity, including gap junctions, plasmodesmata, and septal pores. SppA possesses C2 domain, which has been implicated in anchoring to plasmodesmata^53^. The SppC CUE domain, possesses conserved ubiquitin-binding MFP and LL motifs^47,48^ (Supplementary Fig. 11). The gap-junction protein Cx43 undergoes ubiquitin-mediated degradation upon abiotic stress^54^, suggesting that ubiquitination could be the shared biochemical process regulating cell-to-cell connectivity in fungi and animals.

Cell-to-cell connectivity via gap junctions^54^, plasmodesmata^55^, and septal pores^10^ can be blocked in response to both biotic and abiotic stresses. In this study, the status of cell-to-cell connectivity was quantitively analyzed in response to cold stress (Fig. 5b). Strain lacking SppA showed an unrestricted cell-to-cell transfer of Dendra2 even under cold stress due to defective septum morphology (Fig. 5b). Three SPP proteins (SppB, SppC, and SppE) had a function in septal pore plugging under both wounding and cold stress (Figs. 4b and 5b), suggesting a common stress-response machinery. In general, SPP proteins responded less to cold stress than to hyphal wounding (Figs. 4b and 5b), suggesting their predominant involvement in protecting flanking cells from wound-caused cytoplasmic loss.

According to the orthologous group and phylogenetic analyses, 13 SPP proteins have orthologs outside of Pezizomycotina (Fig. 7). Particularly, in the SPP proteins with orthologs from early diverging fungi, their origins predate the emergence of septal pore, suggesting co-option of the preexisting genes for septal pore regulation. Our bioinformatics-based screening was originally designed to identify genes that were divergent or absent in septal pore–lacking yeasts (Fig. 1b). Contrary to our aim, 12 SPP proteins showed orthologous relations with those of the septal pore–lacking species from ascomycete yeasts and early diverging fungi (Fig. 7, Supplementary Fig. 8). This finding raises a fundamental question regarding the principal and general functions of these proteins. SppA exhibits an extension at the C-terminus in Pezizomycotina (Fig. 8a-d), which could be evolutionarily specific to the porous septum formation. Yeast cytokinesis is morphologically different from that of Pezizomycotina, as yeast daughter cells are completely separated from the mother by SppA orthologs^29,30^. In this scenario, the inability of the sppA deletion mutant to plug the septal pore can be explained by defective cytokinesis or septum formation. Nif1 and Dsf2 are orthologs of SppD in fission and budding yeasts, respectively (Supplementary Fig. 8d). Nif1 acts as a mitotic inhibitor via the interaction with Nim1 protein kinase56, whereas Dsf2 localizes to the bud neck57, the site of cytokinesis. This suggests a possible role of SppD in cell cycle regulation or cytokinesis. SppD subclade is distinct from another subclade containing Pezizomycotina and yeast orthologs (Supplementary Fig. 8d). Therefore, we hypothesize that SppD evolved in Pezizomycotina by gene duplication for performing septal pore-related function. Similarly, fission yeast aim21 is an ortholog of SppV (Supplementary Fig. 8k) and shows dynamic protein turnover during cell division58. Functional roles of SppD and SppV in porous septum formation require further investigation.

SppC possesses a functionally important N-terminal region containing the CUE domain, which shows diversity of the domain presence or absence among its orthologs (Fig. 8e-g, Supplementary Figs. 8b and 11). The Cue2 protein, SppC ortholog in budding yeast, plays a role in ubiquitin binding48. SppJ containing DnaJ domain exhibits an orthologous relation with budding yeast Sis1 (Supplementary Fig. 8e), which is involved in the degradation of cytosolic misfolded proteins59. However, the involvement of ubiquitination and misfolded protein responses in septal pore regulation remains to be investigated. In contrast, SppB does not possess any orthologs in ascomycete yeasts (Fig. 7). The transglutaminase domain of SppB is known for crosslinking properties, which function in mammalian wound healing and blood clotting upon injury^44^. This function is similar to wound-induced plugging of the fungal septal pore.

In the fungal kingdom, hyphal morphology is supposed to have evolved in early diverging fungal phyla Blastocladiomycota, Chytridiomycota, and Zoopagomycota^20^. Septal pore and its regulation appear to have emerged independently in the subphyla Pezizomycotina and Agaricomycotina^1^. Therefore, during the evolutionary transition from aseptate ancestors, many gene evolutionary events were needed to modulate septal-pore organization and regulation. Our orthologous group and phylogenetic analyses (Fig. 7 and Supplementary Fig. 9) demonstrated that 10 SPP genes are present specifically in Pezizomycotina. Such lineage-specific genes have been proposed to have arisen either from ancestrally non-genic regions or by excessive divergence following genomic rearrangement such as recombination, retrotransposition, and horizontal gene transfer^60,61^. The specific existence of the 10 SPP proteins within Pezizomycotina (Fig. 7 and Supplementary Fig. 9) suggests their emergence in unknown ancestor and subsequent acquisition by vertical transfer for aiding septal pore–related functions. This phenomenon may be similar to that underlying multiple lineage-specifically genes present in yeast and worms that are functionally associated with chromosomal segregation^63^.

Our findings show the effectiveness of bioinformatics-based selection to identify many novel proteins involved in fungal septal pore plugging upon hyphal wounding. During the evolution of the septal structure, the interconnected cellular organization might have been vulnerable to environmental threats and physiological stresses such as mechanical wounding^12^, uninterrupted spreading of mycoviruses^63^, and allorecognition-mediated heterokaryon incompatibility^64^. Such adverse conditions could impose selection pressure for adaptation either by the co-option of preexisting genes or by the acquisition of new genes for aiding in septal pore regulation.

## METHODS

### Selection of candidate septal pore–related proteins

Whole proteomes of seven fungal species were retrieved from the genome database FungiDB (https://www.fungidb.org) and subsequent assemblies. We used 12090 *A. oryzae* proteins to query each using BLASTp and mpiBLAST. The presence or absence of the septal pore, as a specialized morphological structure, was set as the criteria for genomic comparison. We searched for conserved genes in septal pore–bearing ascomycetes *A. oryzae*, *A. fumigatus,* and *A. nidulans* with e-values less than or equal to 1e-100; we then searched for genes absent or diverged in the ascomycete yeasts *S. cerevisiae*, *S. pombe,* and *C. albicans*, with e-values greater than or equal to 1e-30. This process yielded 2130 candidates for septal pore-related proteins in *A. oryzae*.

We preferentially selected uncharacterized proteins, defined as those lacking biological data for GO terms regarding molecular functions, cellular components, and biological processes. A total of 5878, 7868, and 5664 genes were identified for molecular functions, cellular components, and biological processes, respectively. A total of 3498 genes lacking biological data for all three GO terms were finally determined as uncharacterized genes. We then searched for the aforementioned 2130 candidate proteins within this uncharacterized subset, and shortlisted 776 genes for localization screening.

### Strains, growth conditions, and transformation

Strains used in this study are listed in Supplementary Data 3. *A. oryzae* was transformed as previously described^65^. *A. oryzae* strain NSPlD1 (*niaD*^−^ *sC*^−^ Δ*pyrG* Δ*ligD*)^10^ was used as the parent strain. Using the *pyrG* selectable marker, transformants were selected using M+Met medium [0.2% NH_4_Cl, 0.1% (NH_4_)_2_SO_4_, 0.05% KCl, 0.05% NaCl, 0.1% KH_2_PO_4_, 0.05% MgSO_4_ ·7H_2_O, 0.002% FeSO_4_·7H_2_O, 0.15% methionine, and 2% glucose; pH 5.5]. Transformants with the *niaD* and *sC* selectable markers were selected using CD medium (0.3% NaNO_3_, 0.2% KCl, 0.1% KH_2_PO_4_, 0.05% MgSO_4_·7H_2_O, 0.002% FeSO_4_·7H_2_O, and 2% glucose; pH 5.5) and CD+Met medium supplemented with 0.0015% methionine, respectively. *A. oryzae* strains were maintained in Potato Dextrose (PD) medium (Nissui, Tokyo, Japan) at 30°C. For transformation, strains were inoculated into DPY liquid medium (0.5% yeast extract, 1% Hipolypeptone, 2% dextrin, 0.5% KH_2_PO_4_, 0.05% MgSO_4_·7H_2_O, pH 5.5) and grown overnight. For microscopy, conidia were inoculated on glass base dishes containing 100 μL CD liquid medium supplemented with 1% casamino acids. For hyphal wounding, agar medium instead of liquid medium was used. For complete induction and repression of the *amyB* promoter, 2% dextrin and 2% glycerol, respectively, were included in CD medium as carbon sources.

### DNA techniques

Nucleic-acid sequences of the 776 candidates septal pore proteins were retrieved from the Aspergillus genome database AspGD (http://www.aspgd.org). The open reading frame encoding each protein was amplified from RIB40 genomic DNA with PrimeSTAR^®^ HS DNA Polymerase (TaKaRa Bio, Otsu, Japan) for high-fidelity PCR, using primers listed in Supplementary Data 6. The amplified fragments were then fused with *Sma*I-linearized pUt-C-EGFP^10^ using the In-Fusion HD Cloning Kit (Clontech Laboratories, Mountain View, CA, USA). Gene sequences fused with the *egfp* gene were positioned in tandem between the linker sequences. Recombinant plasmids used for the expression of EGFP-fused proteins were digested with *Not*I and inserted into the *niaD* locus of the wild-type strain NSlD1^10^ by homologous recombination. Transformants were selected by nitrate assimilation. EGFP fusion–expressing strains generated for localization screen are listed in Supplementary Data 3.

Primers used for the generation of the deletion strains are listed in Supplementary Data 6. Individual genes with the *pyrG* marker were replaced by amplifying the 1.5 kbp regions upstream and downstream of the genes using the primer sets gene-upstream1_F/gene-upstream2_R and gene-downstream1_F/gene-downstream2_R, respectively. The *pyrG* marker was amplified from RIB40 genomic DNA using the primers PyrG_F and PyrG_R. The three amplified DNA fragments (upstream and downstream regions of targeted genes and *pyrG*) and linearized-pUC19 vector were fused using the In-Fusion HD Cloning Kit. Deletion constructs were further amplified using the primer set gene-upstream1_F/gene-downstream2_R from the resultant plasmid as template, and then introduced into the native locus of the *A. oryzae* strain NSPlD1^10^ by homologous recombination. Transformants were selected using uridine/uracil prototrophy.

Deletions of *sppG*, *sppF*, and *sppE* with the *sC* marker were performed by amplifying 1.5 kbp upstream and downstream regions using the primer sets gene_upstream1_F/gene-upstream2 (sC)_R and gene-downstream1(sC)_F/gene-downstream2_R, respectively. The *sC* marker was amplified from RIB40 genomic DNA using the primers sC_F and sC_R. The three fragments and linearized-pUC19 vector were fused. PCR-amplified deletion constructs of *sppG*, *sppF*, and *sppE* were introduced into the native loci of Δ*sppA*/Δ*sppP*/Δ*sppI*/Δ*sppT*/Δ*sppW*/Δ*sppL*/Δ*sppS*/Δ*sppJ*/Δ*sppQ*, Δ*sppM*/Δ*sppH*/Δ*sppO*/Δ*sppU*/Δ*sppR*, and Δ*sppB*/Δ*sppC*/Δ*sppD*/Δ*sppK*/Δ*sppN*/Δ*sppV*, respectively.

To analyze cell-to-cell connectivity via the septal pore under cold stress (Fig. 5), Dendra2, a green-to-red photoconvertible fluorescent protein, was used^10^. The Dendra2–expressing plasmid pUt-NA-*dendra*2^10^ was amplified and digested with *Not*I. Ethanol-precipitated DNA was inserted into the *niaD* locus of deletion strains lacking 62 septum-localizing proteins by homologous recombination.

To determine the importance of the disordered region, fragments encoding residues 1-264 and 265-697 of SppB; 1-145 and 146-390 of SppD; 1-265 and 265-1234 of SppN; 1-92, 93-509, 1-192 and 193-470 of SppI; 1-99 and 100-962 of SppL; and 1-167 and 168-465 of SppW were amplified. For the truncated variants lacking the N-terminal region, a start codon was added. The individual fragment was then fused with *Sma*I-linearized pUt-C-EGFP. Recombinant plasmids were digested with *Not*I and inserted into the *niaD* locus of the corresponding deletion backgrounds by homologous recombination.

The primers used to truncate SppA and SppC are listed in Supplementary Data 6. The amplified truncated fragments were fused with *Sma*I-digested pUt-C-EGFP, and then a *Not*I-digested expression cassette was inserted into the *niaD* locus. To generate chimeric proteins of SppA (Fig. 7), a DNA encoding the N-terminus (residues 1-127) was amplified from RIB40 genomic DNA, and those for the C-termini of *A. nidulans* (residues 126-810), *N. crassa* (residues 127-863), and *S. pombe* (residues 135-810) were amplified using the genomic DNA of corresponding strains FGSC_A26, OR74A, and 972h, respectively. The fragments were fused with *Sma*I-digested pUt-C-EGFP^10^ using the In-Fusion HD Cloning Kit. The constructed plasmids were digested with *Not*I and inserted into the *niaD* locus of Δ*sppA* by homologous recombination.

SppC was truncated from both the C- and N-termini. DNA fragments encoding N-terminal residues 1-142 and 1-185 were amplified from RIB40 genomic DNA. They were fused with *Sma*I- digested pUt-C-EGFP. To truncate N-terminal regions containing the CUE domain, a DNA fragment encoding residues 143-556 was amplified by adding an ATG start codon. The fragment and the *Sma*I-digested pUt-C-EGFP were fused using the In-Fusion HD Cloning Kit. The constructed plasmids were digested with *Not*I and used to insert expression cassettes into the *niaD* locus of the Δ*sppC* by homologous recombination.

### Microscopy

Apical septa of living hyphae expressing EGFP-tagged proteins were observed using an IX71 inverted microscope (Olympus, Tokyo, Japan) equipped with 100× Neofluar objective lenses (1.40 numerical aperture); 488-(Furukawa Electric, Tokyo, Japan) and 561-nm (Melles Griot, Rochester, NY, USA) semiconductor lasers; GFP filters (Nippon Roper, Tokyo, Japan); a CSU22 confocal scanning system (Yokogawa Electronics, Tokyo, Japan); and an Andor iXon cooled digital CCD camera (Andor Technology PLC, Belfast, UK). Images were analyzed using Andor iQ 1.9 software (Andor Technology PLC).

### Hypotonic shock and Dendra2 transfer analyses

Hyphal tips were burst by flooding colonies with 1 mL water after growth on a thin layer of DPY agar medium in a glass base dish at 30°C for 18 h. After flooding, the colony was left for 2 min before observation. Thirty randomly selected hyphae showing burst tips were observed using differential interference contrast microscopy.

To analyze the Dendra2 transfer analysis, hyphae were grown on a thin layer of CD (2% dextrin as carbon source) medium supplemented with 1% casamino acids at 30°C for 17 h. The culture was further incubated in normal or cold stress (4°C, 30 min) conditions. Before observation, hyphae were illuminated with a UV mercury lamp positioned at least 20 μm from the selected apical septum to avoid illumination of the adjacent cell, as previously described^10^. Dendra2 transfer via the septal pore was observed after photoconversion with 553-nm excitation^10^.

### Disorder prediction and analysis of amino acid composition

Disordered regions of SPP proteins were annotated using IUPred2A^31^. To analyze relative amino acid composition (Supplementary Fig. 7c), the ordered proteins were obtained from the O_PDB_S25 dataset^66^ (*b*) and the disordered set was retrieved from the DisProt database^67^. The serine/ arginine (SR) and phenylalanine/glycine (FG) datasets were previously defined^68,69^. The sequences of SPA proteins were retrieved from the FungiDB genome database. The disorder regions (*a*) were taken from SPP and SPA using IUPred2A, as well as DisProt, SR, and FG datasets. The fractional difference was calculated^70^ by (*M ^a^* amino acid. - *M ^b^*) /*M_j_ ^b^*, where *j* is for the individual

### Orthologous group and phylogenetic analyses

Twenty-three SPP proteins were analyzed using Simple Modular Architecture Research Tools (SMART: http://smart.embl-heide lberg.de/)^66^ to detect predicted domains. To analyze primary sequence conservation, multiple sequence alignments were performed using ClustalW^67^, and outputs were presented using BoxShade server version 3.21.

For orthologous group analysis, we obtained fungal proteome data available from the National Center for Biotechnology Information (NCBI) (https://www.ncbi.nlm.nih.gov/) and MycoCosm of Joint Genome Institute (JGI) fungal portal (https://mycocosm.jgi.doe.gov/mycocosm/home) databases. In total, 81 fungal species were selected to cover all major clades of fungi from subphyla of Ascomycota (Taphrinomycotina, Saccharomycotina, and Pezizomycotina) and Basidiomycota (Agaricomycotina, Pucciniomycotina, and Ustilaginomycotina), as well as representative phyla of Mucoromycota, Zoopagomycota, Chytridiomycota, Blastocladiomycota, and Rozellomycota (Supplementary Data 7). Orthologous groups were analyzed using Orthofinder 2.3.14^35^ with default parameters.

For substitution rate analysis, we followed a previously described method^21^. A representative protein that phylogenetically close to an orthologous Pezizomycotina protein was selected from each fungal species (Supplementary Data 5d). Multiple sequence alignments were built for individual proteins using ClustalW^68^ with unified gap penalties, including gap open penalties and gap extension penalties. Maximum likelihood trees were generated for each protein using PhyML. Phylogenetic trees were confirmed by constructing them with a different tool, Molecular Evolutionary Genetics Tool Mega^71^. The substitution rate was calculated for a given species, *x*, in the corresponding tree, and the evolutionary distance between protein sequences and Pezizomycotina ancestral sequences of A. oryzae SPP proteins was estimated by the score 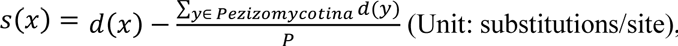 where *d*(*x*) indicates the branch length from species *x* to the Pezizomycotina root. *P* represents the number of species *y* in Pezizomycotina used in phylogenetic analysis, and therefore 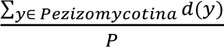 represents the average branch length from each species *y* in Pezizomycotina to the Pezizomycotina root.

To further elucidate orthologous relationships, phylogenetic trees were generated using all proteins classified into the same orthologous group. Multiple sequence alignments were performed and unrooted phylogenetic trees for analyzed proteins were constructed using Mega^71^.

### Western blot analysis

Strains were cultured on DPY medium for 18 h. The fungal mycelia were frozen in liquid nitrogen and subsequently ground with multi-bead shocker. After extracting, cell lysates were incubated with the buffer containing detergent NP-40 (50 mM Tris-HCl (pH 8.0), 200 mM NaCl, 1 mM/mL PMSF, 1 mM/mL Protease inhibitor, 1% NP-40) to solubilize the membrane protein. Mycelial extract solubilized with the buffer was incubated in ice for 20 min. After centrifugation with 1000x *g* for 10 min, the supernatant was collected. Cell lysates were separated using SDS-PAGE. The proteins were transferred to Immobilon-P polyvinylidene difluoride (PVDF) membranes (Millipore Sigma, Burlington, MA, USA) using a semidry blotting system (Nihon Eido, Tokyo, Japan). To detect EGFP, Living Colors A.V. monoclonal antibody (Clontech) and peroxidase-labeled anti-mouse IgG (H + L) antibody (Vector Laboratories, Newark, CA, USA) were used as primary and secondary antibodies, respectively. Chemiluminescence was detected using a Western Lightning-ECL system (PerkinElmer, Waltham, MA, USA) and an LAS-4000 image analyzer (GE Healthcare, Buckinghamshire, UK).

### Statistical analysis

The results of at least three independent experiments are presented as means. Error bars represent standard deviations as indicated in the figure legends. Statistical significance was tested with the two-tailed Student’s *t-*test using Microsoft Excel; significance is indicated as *\*p* < 0.05 or *\*\* p* < 0.01.

### Data availability

All data supporting the findings of the present study are available in this article and Supplementary Information files, or from the corresponding authors upon request.

## ACKNOWLEDGEMENTS

This study was supported by Japan Society for the Promotion of Science (JSPS) KAKENHI Grant Number 21H02098, the Grant-in-Aid for Scientific Research (B) and 21F21099, and the Grant-in-Aid for JSPS Fellows.

## AUTHOR CONTRIBUTIONS

M.A.A.M and J.M. conceived of and designed the study. M.A.A.M. performed experiments, and M.A.A.M. and J.M. interpreted the results. S.N. and W.C. carried out the large-scale BLAST search and interpreted the data. J.M. performed orthologous group and phylogenetic analyses, and all authors interpreted the data. M.A.A.M and J.M. wrote the manuscript. and approved of the final manuscript. All authors read and approved the manuscript.

**COMPETING INTERESTS**

The authors declare no competing interests.

---

**Supplementary Figure 1:**
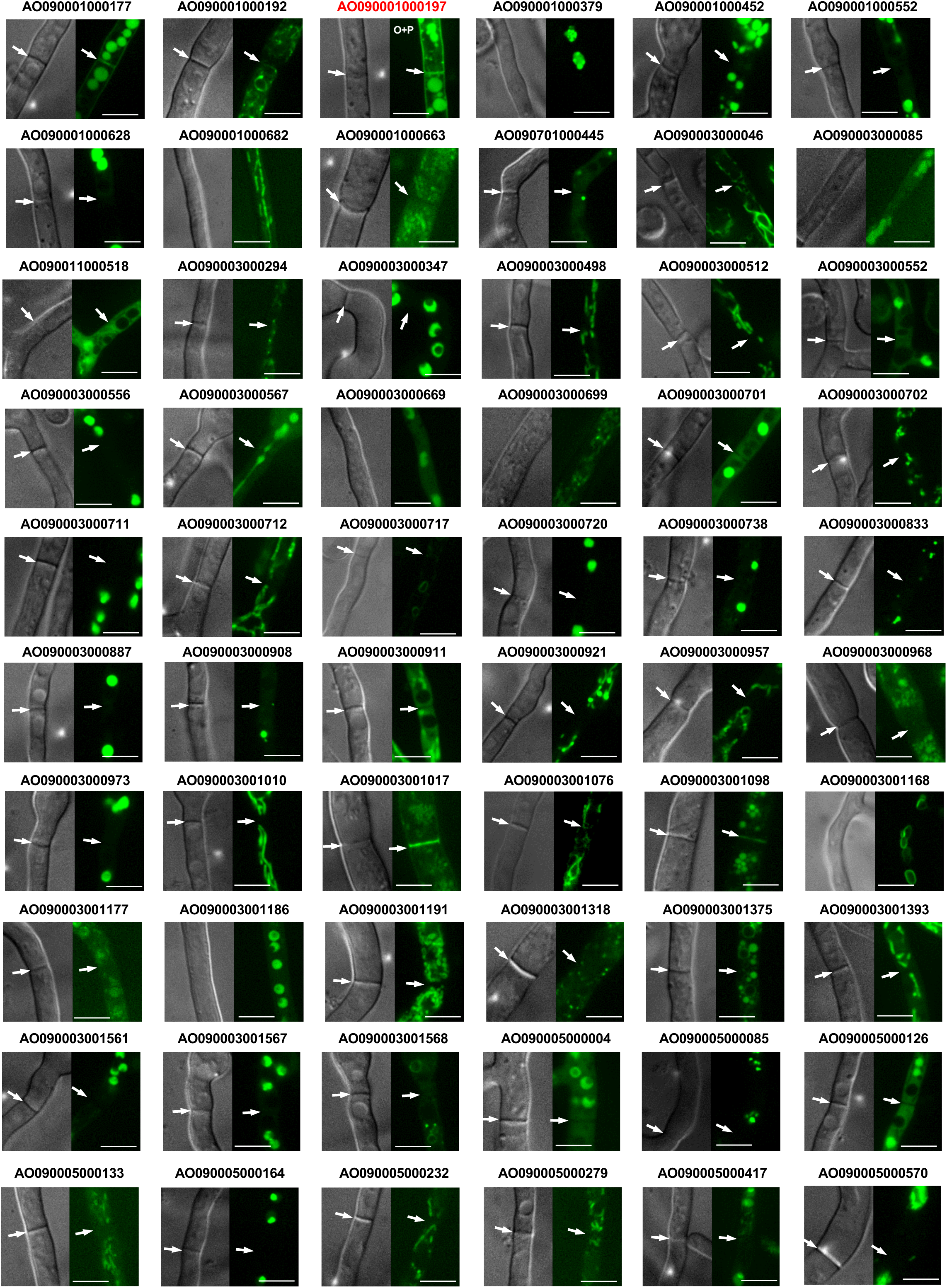
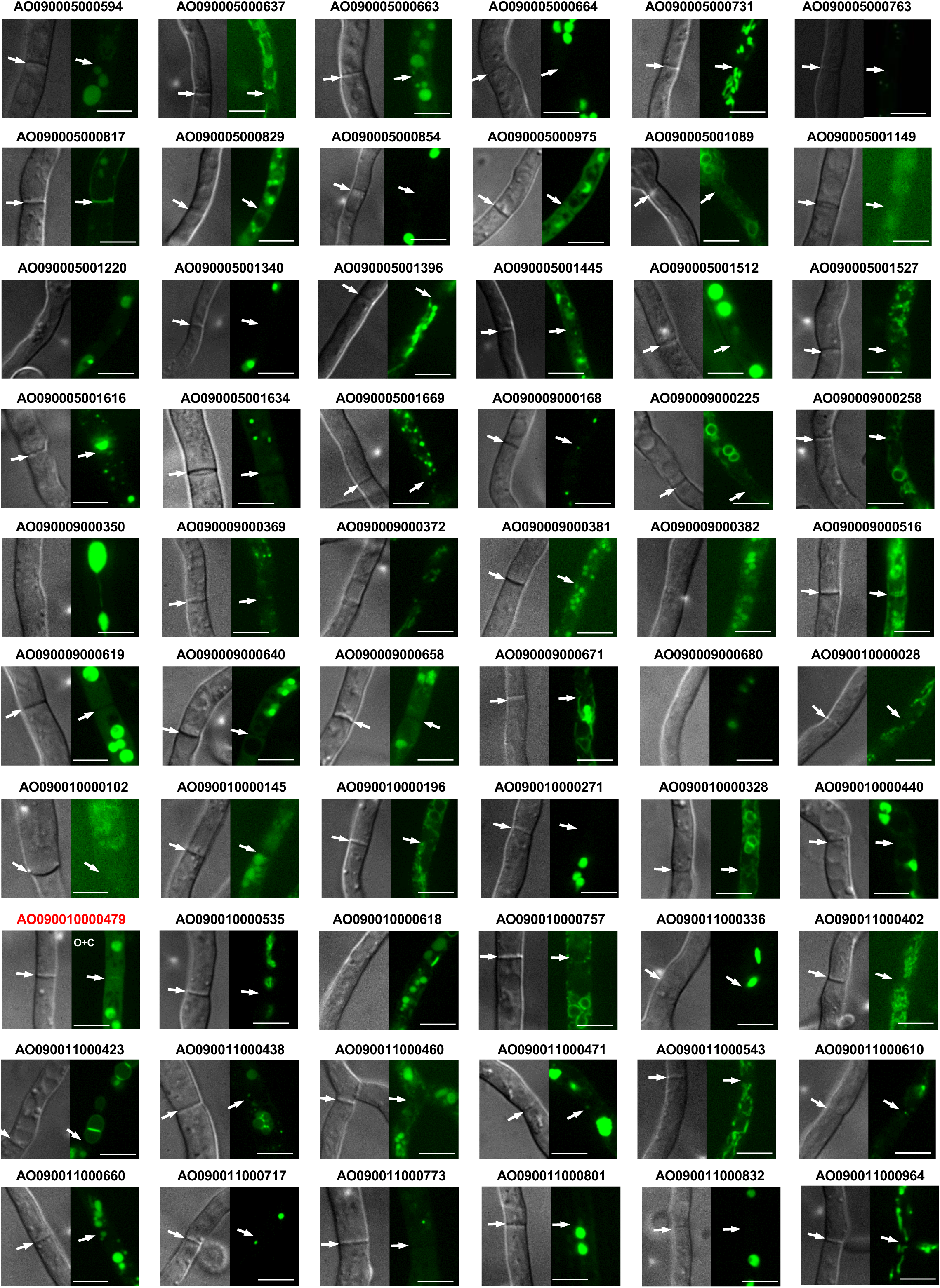
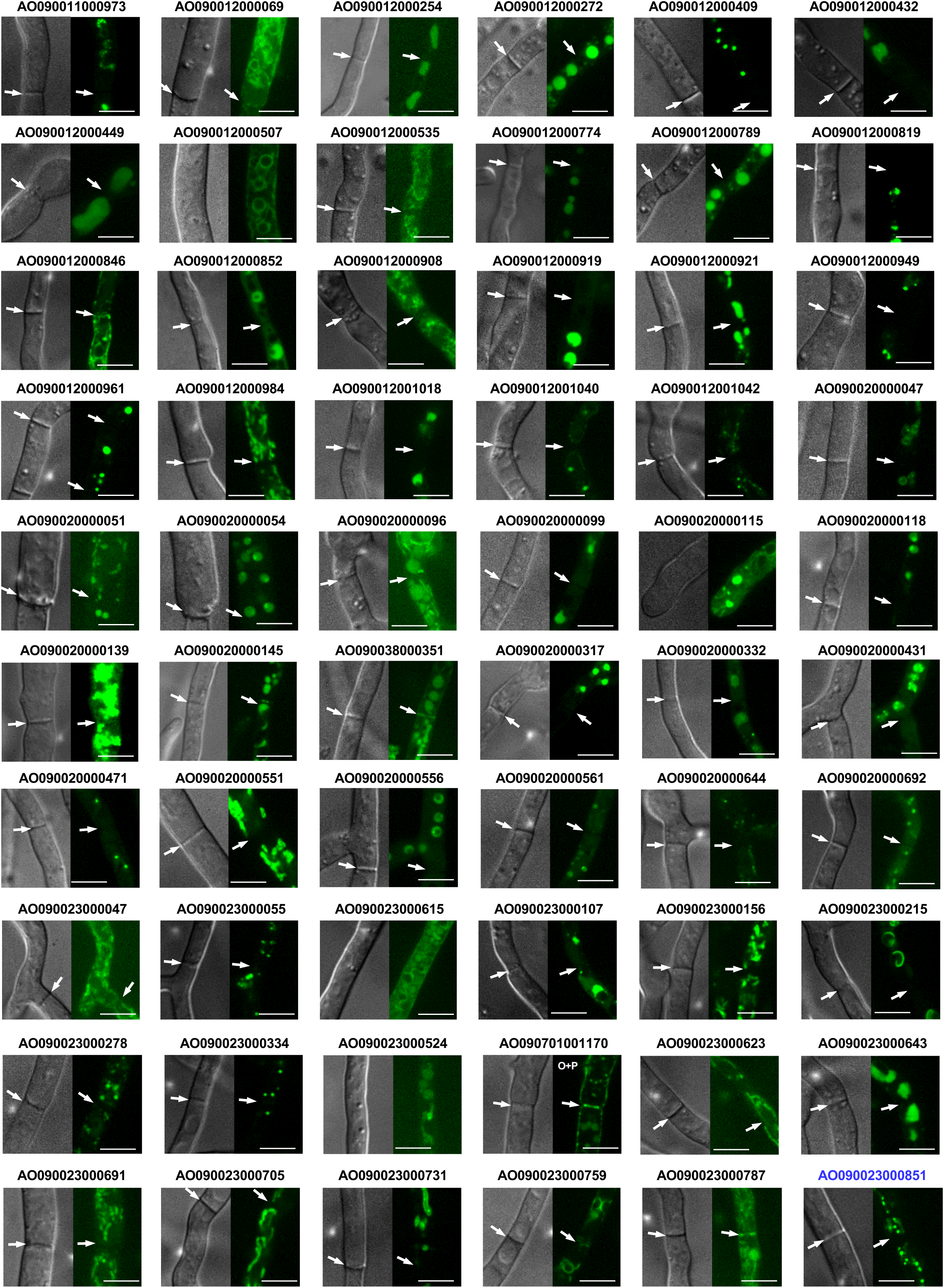
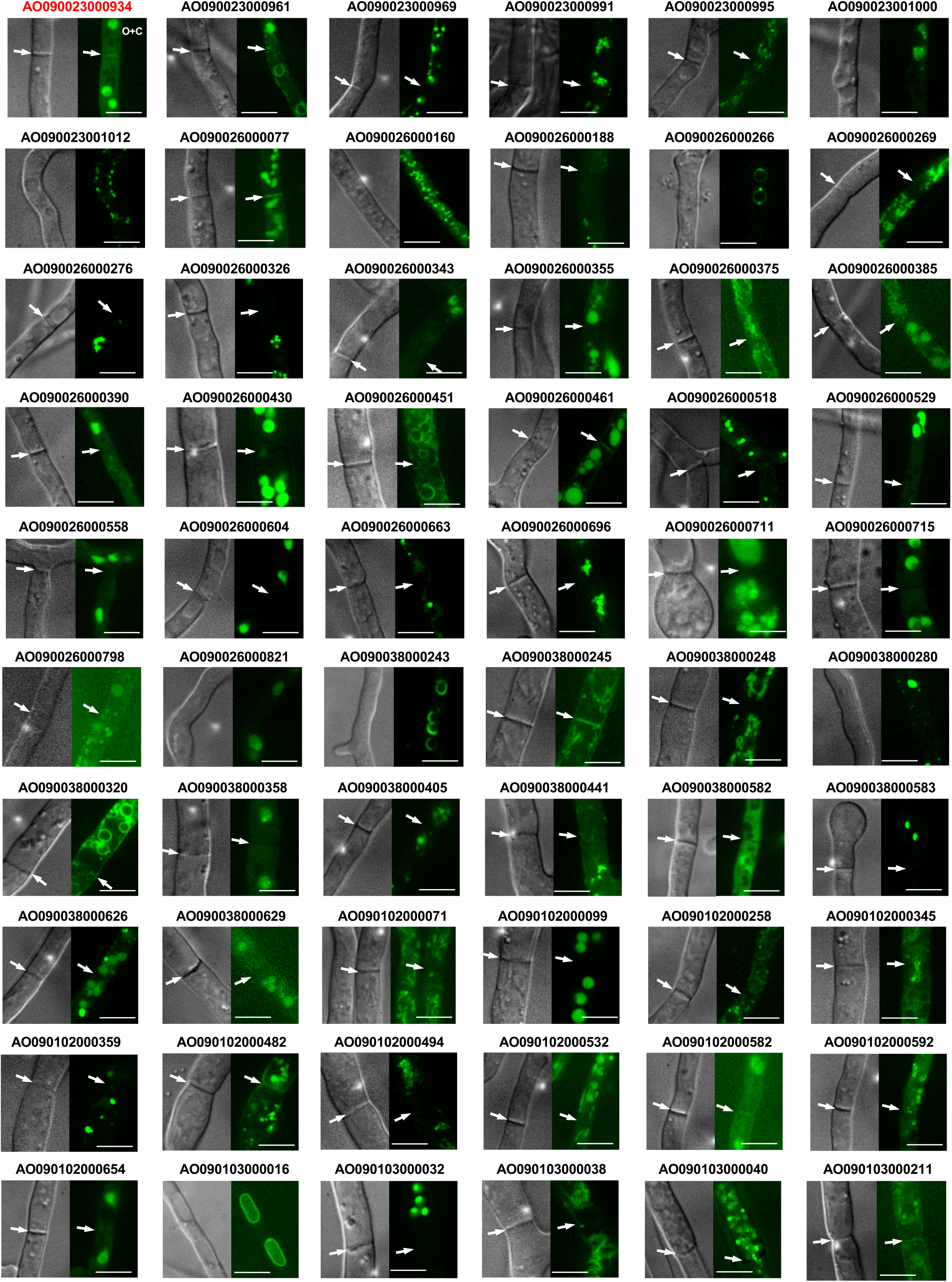
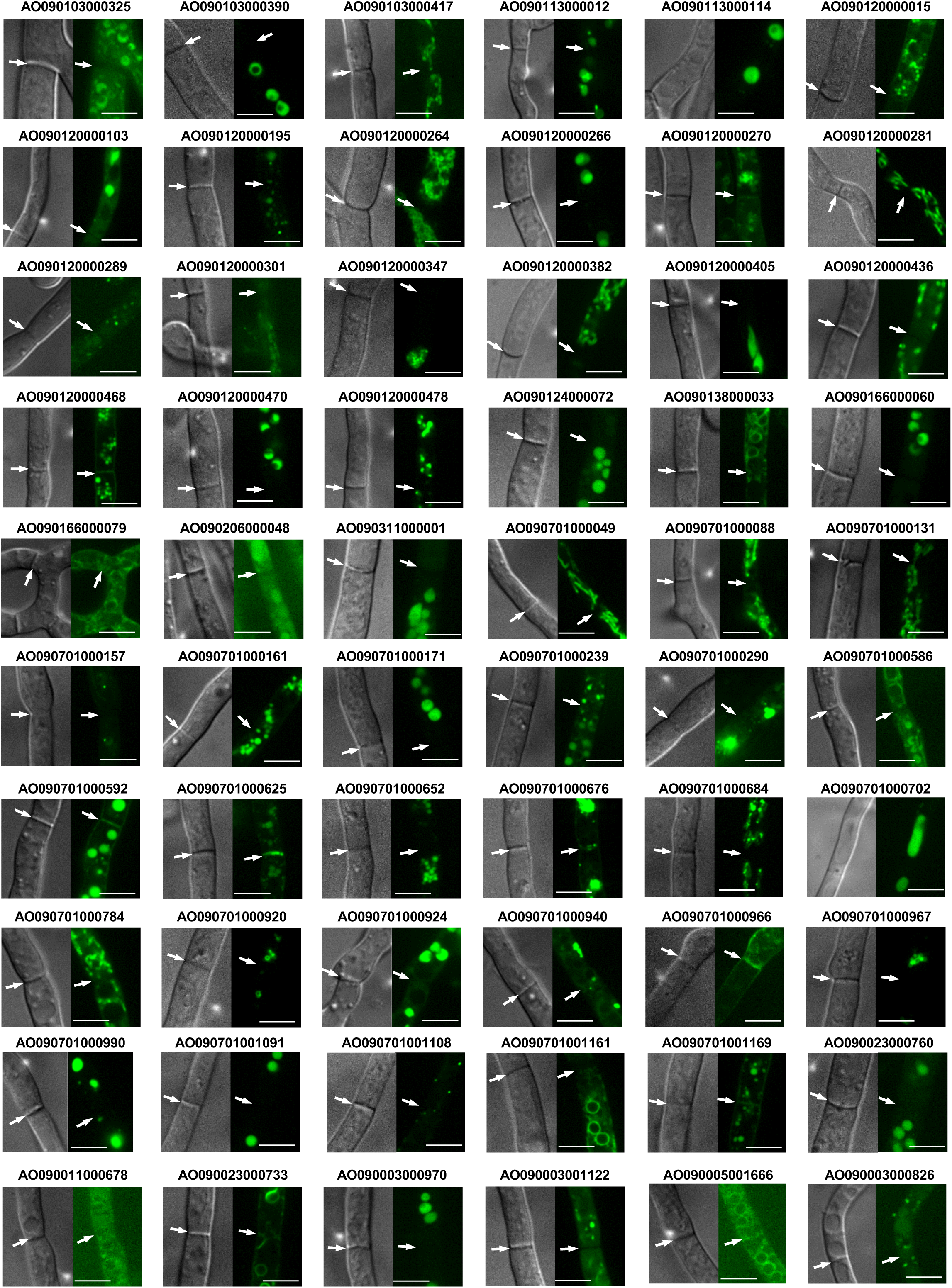
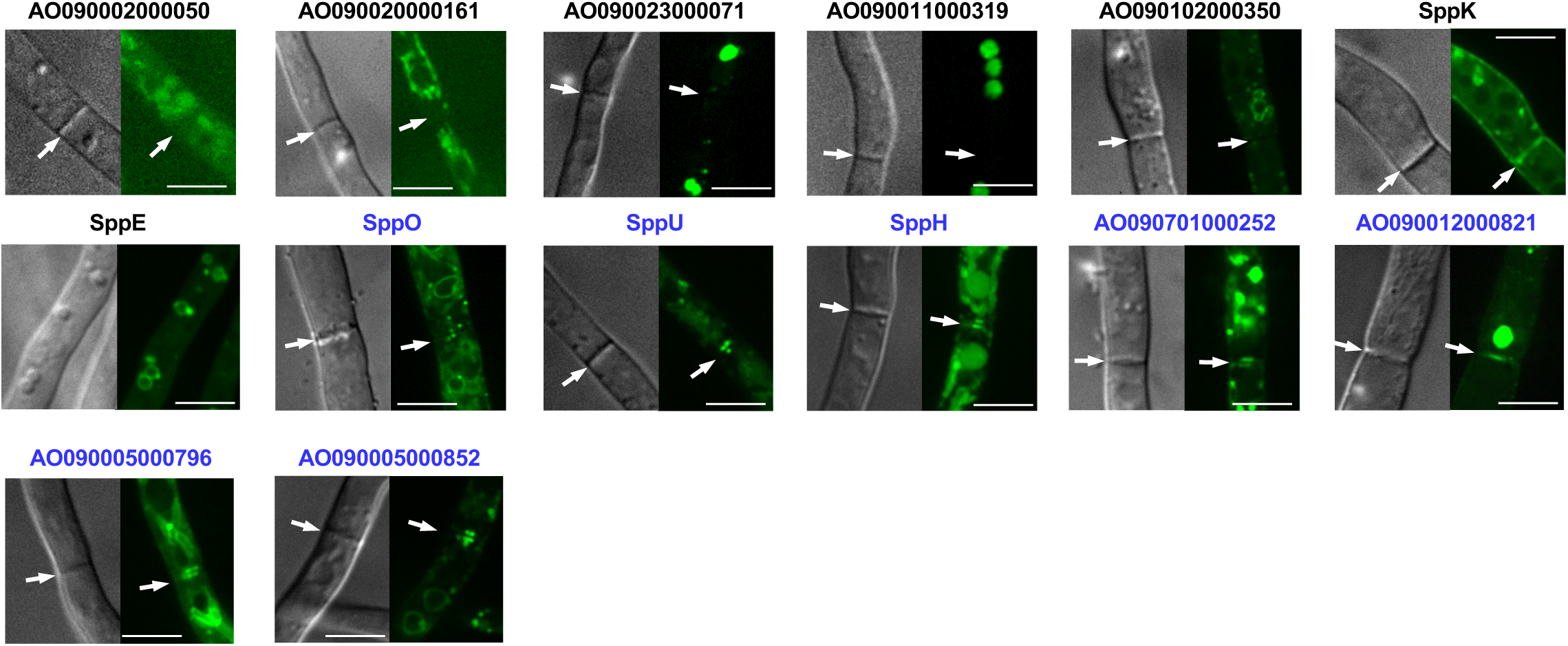
Organelle-like localization of 314 candidate proteins. Panel labels are Gene IDs or SPP protein names, and the proteins, which localized to the septum, are shown in blue. Other overlapped localizing proteins were indicated in red. Arrows indicate septa. Scale bars, 5 μm.

**Supplementary Figure 2:**
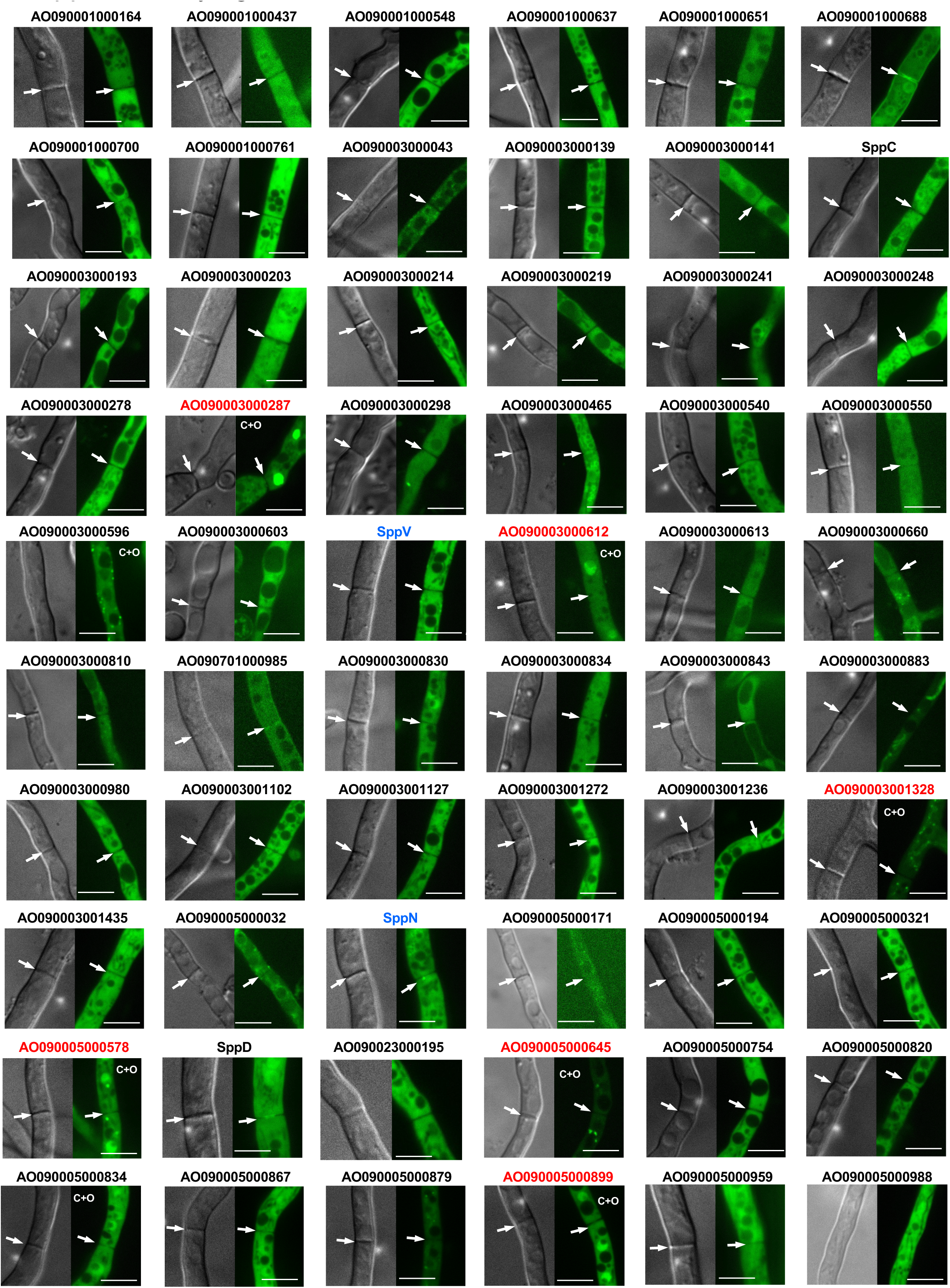
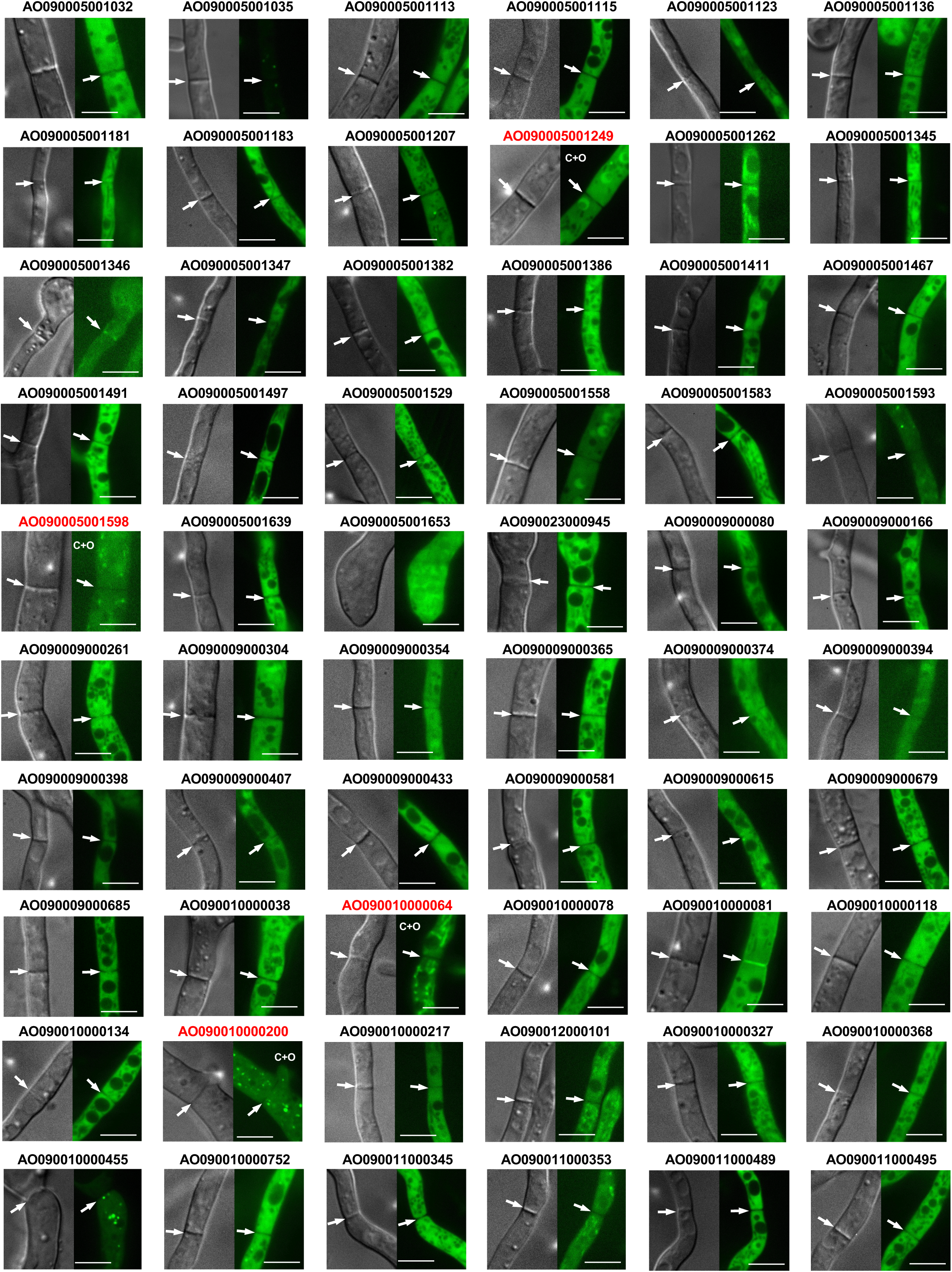
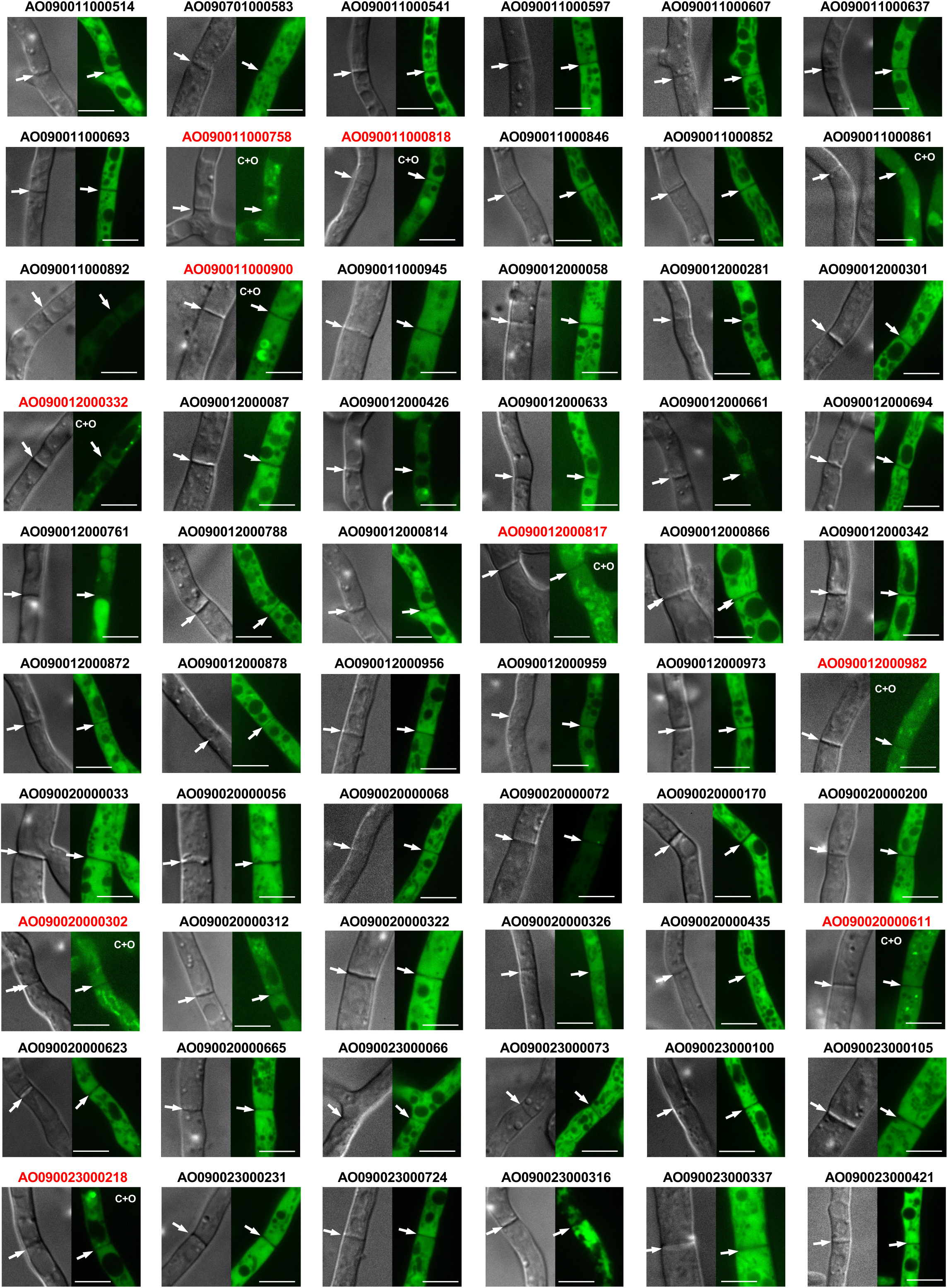
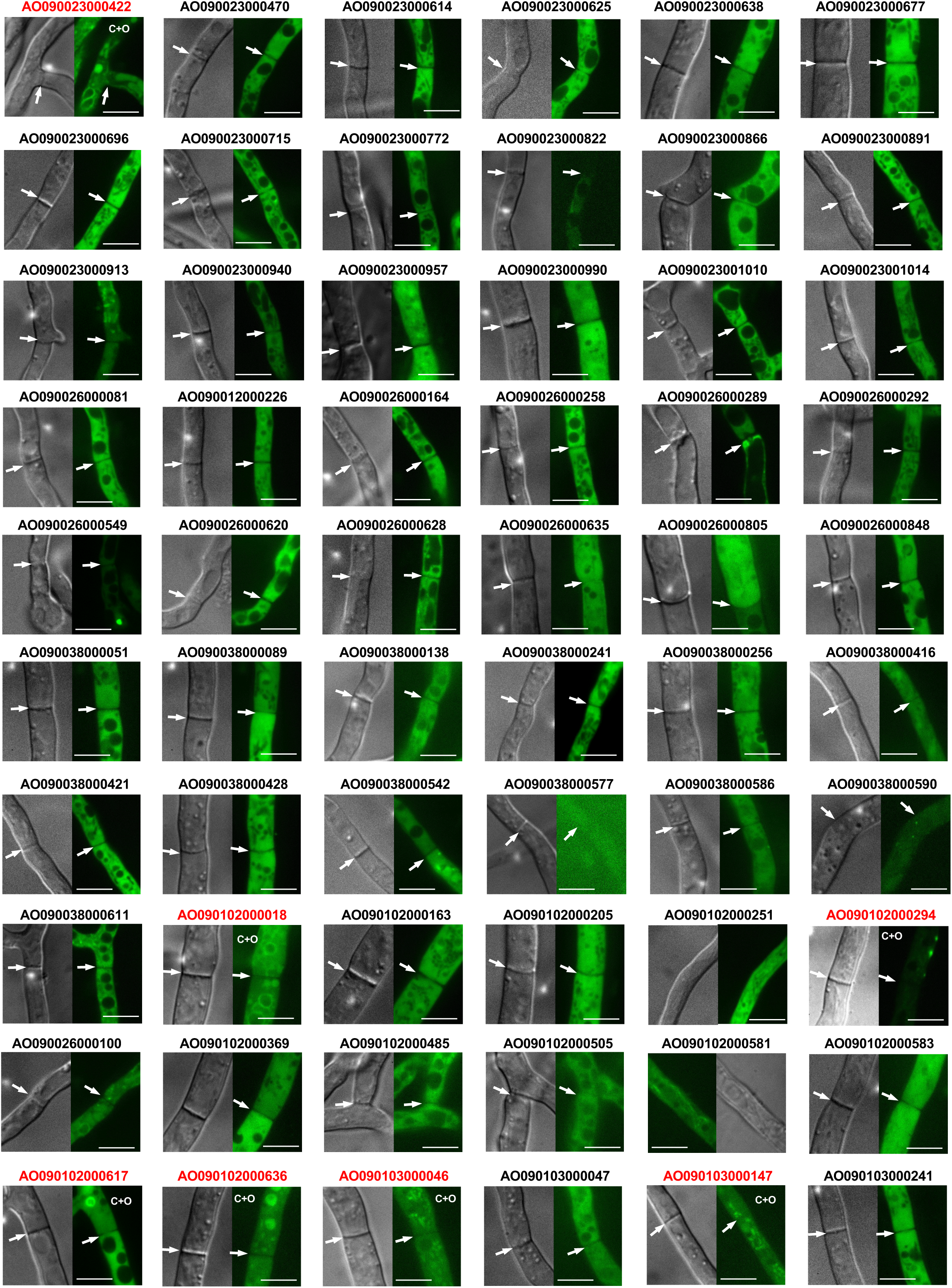
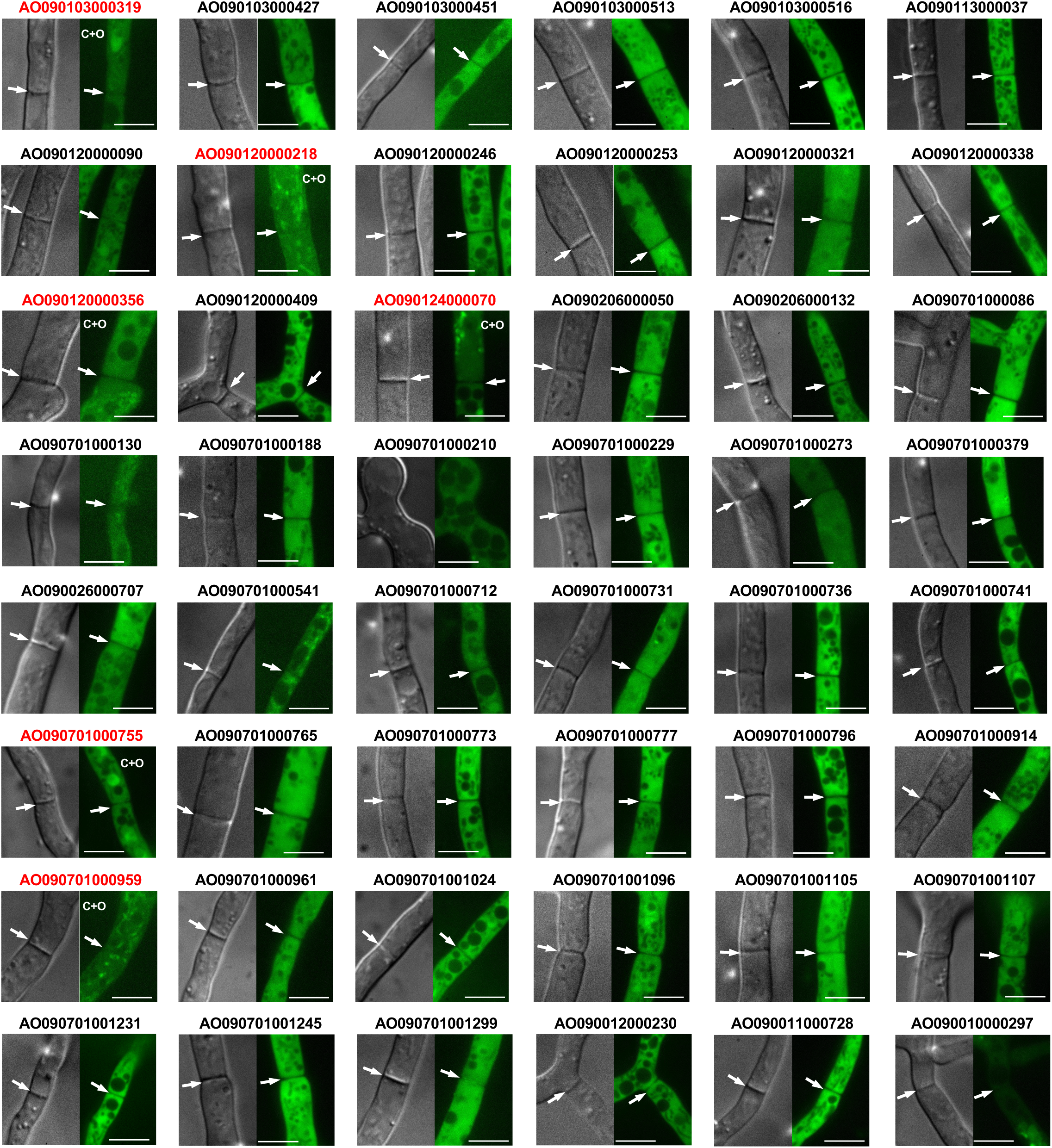
Cytoplasmic localization of 288 candidate proteins. Gene IDs for the proteins are represented for the corresponding micrographs, and the proteins, which localized to the septum, are shown in blue. Arrows indicate septa. Scale bars, 5 μm.

**Supplementary Figure 3:**
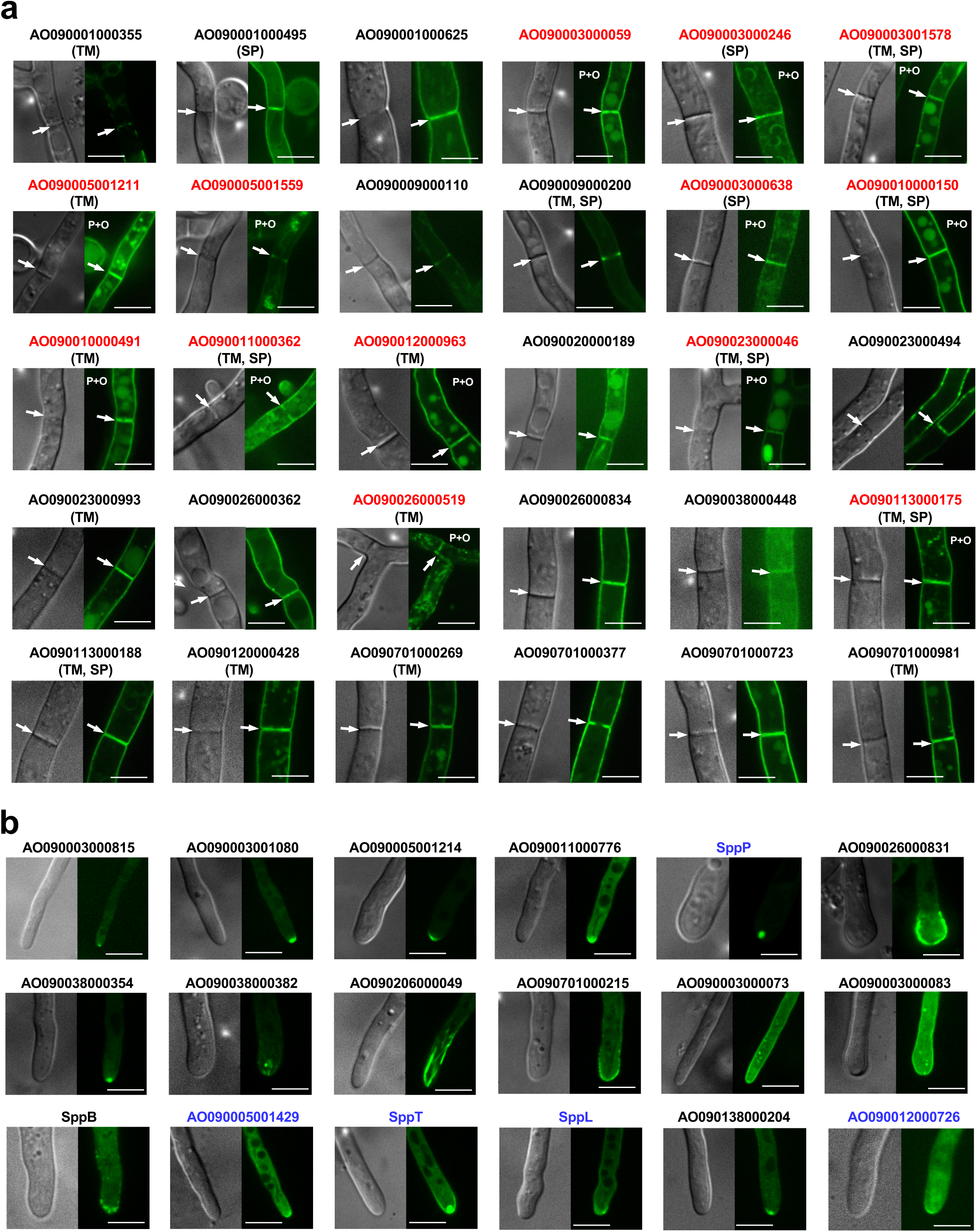
Peripheral and hyphal tip localization of candidate proteins. **a** Peripheral localization of 30 proteins. TM and SP under gene IDs indicate transmembrane domain and signal peptide, respectively. **b** Hypha!tip localization of 18 candidate proteins. The proteins, which localized to the septum under normal growth condition, are shown in blue. Arrows indicate septa. Scale bars, 5 fUll.

**Supplementary Figure 4:**
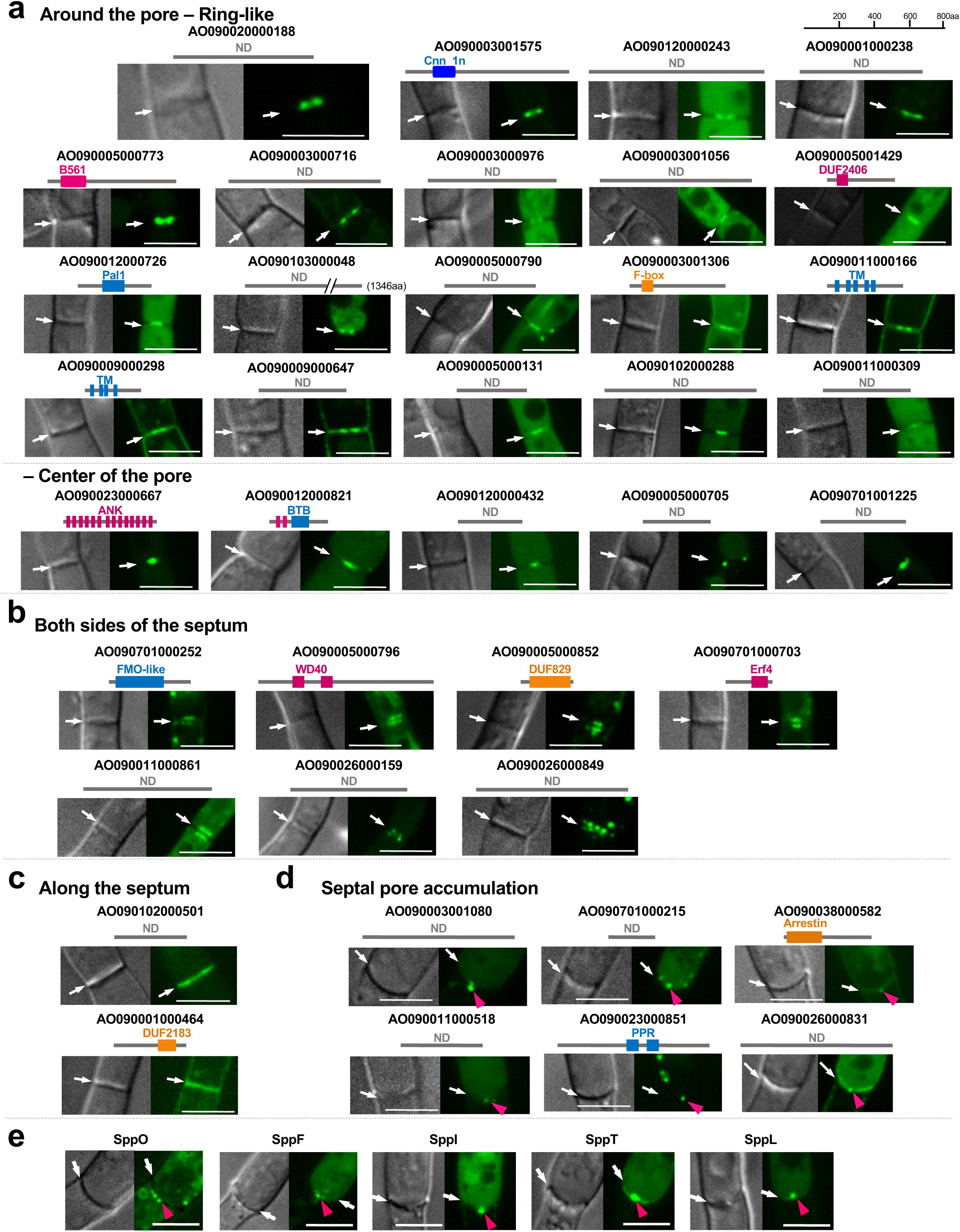
Septum and septal pore localization of candidate proteins. a Protein localizations around the septal pore. **b** Protein localizations on both sides of the septum. c Proteins localizations along the septum. **d** Accumulation of non-septal proteins at the septal pore upon hyphal wounding. e Septal pore accumulation of proteins, which localized to the septum under normal growth condition, upon hyphal wounding. Arrows indicate septa, and arrowheads represent septal pore accumulation. Scale bars, 5 fllTI. Cartoons illustrate the pattern of localizations at the septum. ND; no predicted domains.

**Supplementary Figure 5:**
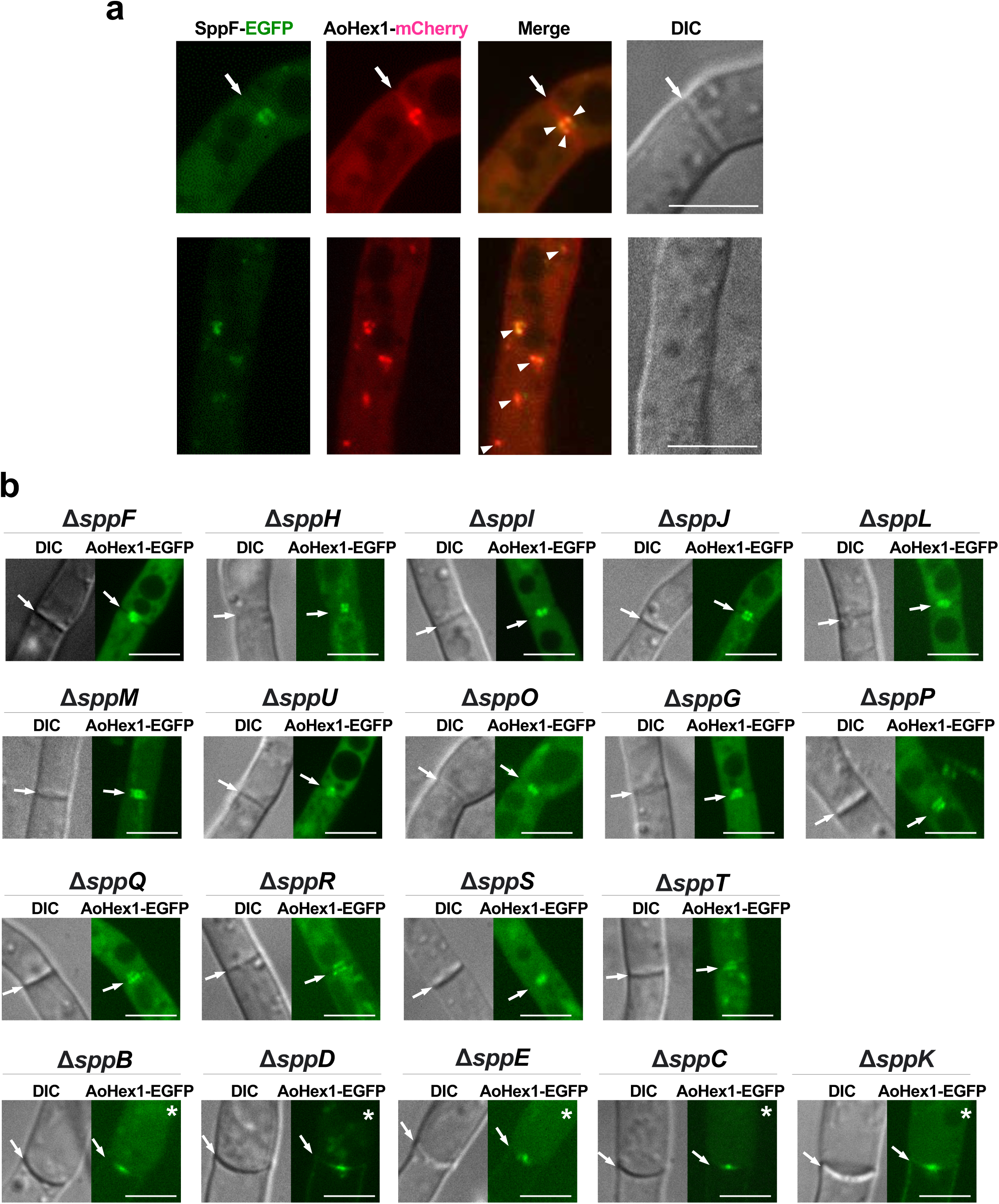
Relationships between SPP proteins and Woronin bodies. **a** Woronin bodies visualized with AoHexl-mCherry and SppF with EGFP. Arrows indicate septa, and arrowheads indicate colocalization. Scale bars, 5 μm. **b** Woronin bodies visualized with AoHexl-EGFP were expressed in *spp* deletion backgrounds. Arrows indicate septa, and asterisks indicate hyphae subjected to hyphal wounding induced by hypotonic shock. Scale bars, 5

**Supplementary Figure 6:**
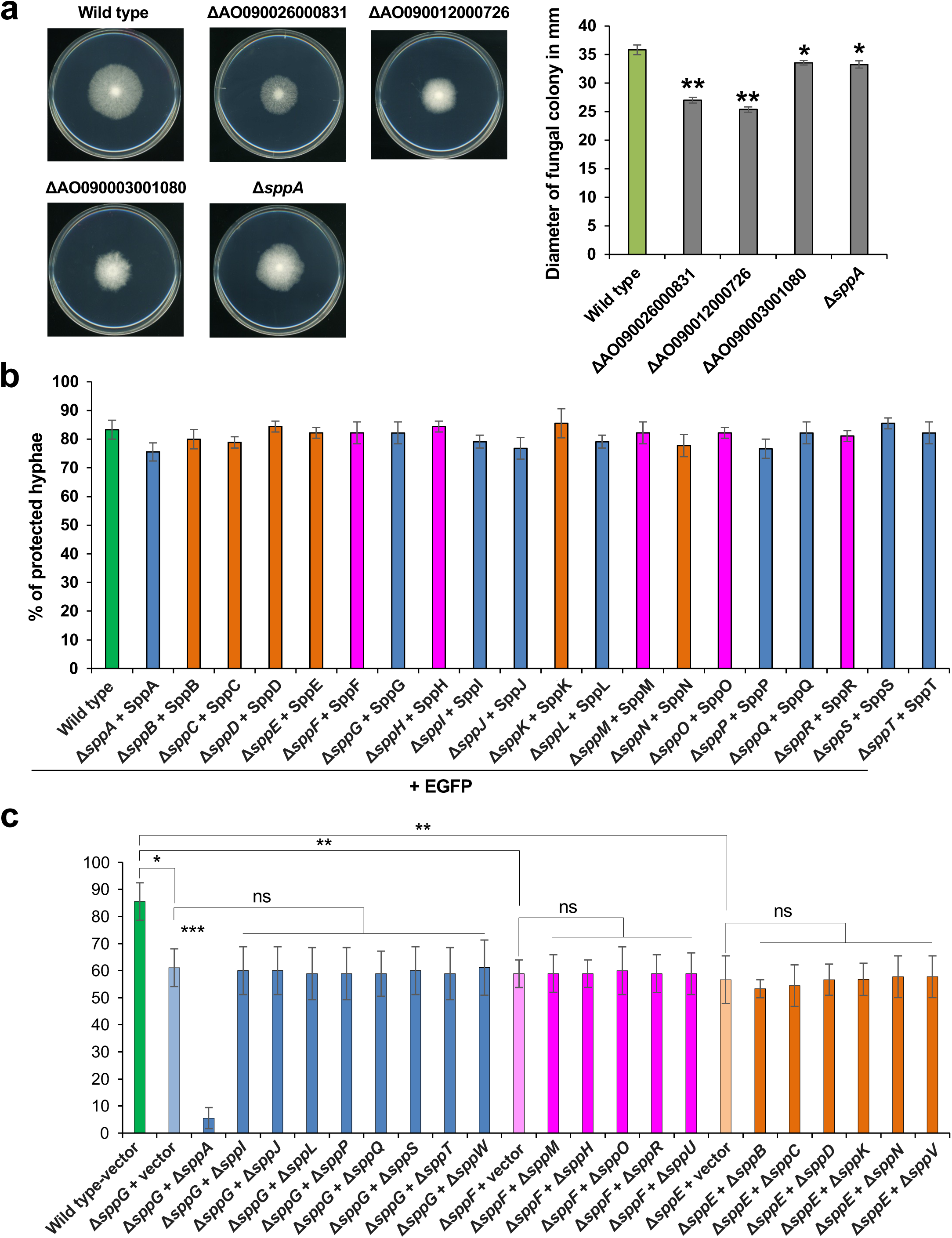
Functionality of EGFP-fused SPP proteins. **a** Colony growth from conidial suspensions. Three independent experiments were performed. Error bars represent standard deviations. *\*p* < 0.05; *\*\*p* < 0.01 (Student’s t-test). **b** Protection of flanking cells from cytoplasmic loss upon hyphal tip bursting by expression of EGFP-tagged SPP proteins on deletion backgrounds. c Protection of flanking cells from cytoplasmic loss upon hyphal tip bursting in the double deletion strains. Thirty randomly selected hyphae showing hyphal tip bursting were observ ed in each experim ent. Three independent experiments were performed, and percentage of hyphae protected from the excessive loss of cytoplasm is shown in the graph. Error bars represent standard deviations. The statistical analysis was performed between the wild type and three representative single deletions representative for each localization category or between the double deletions and the corresponding representative single deletion. *\*p* < 0.05; *\*\*p* < 0. 01; ns; not significant (Student’s t-test).

**Supplementary Fig. 7:**
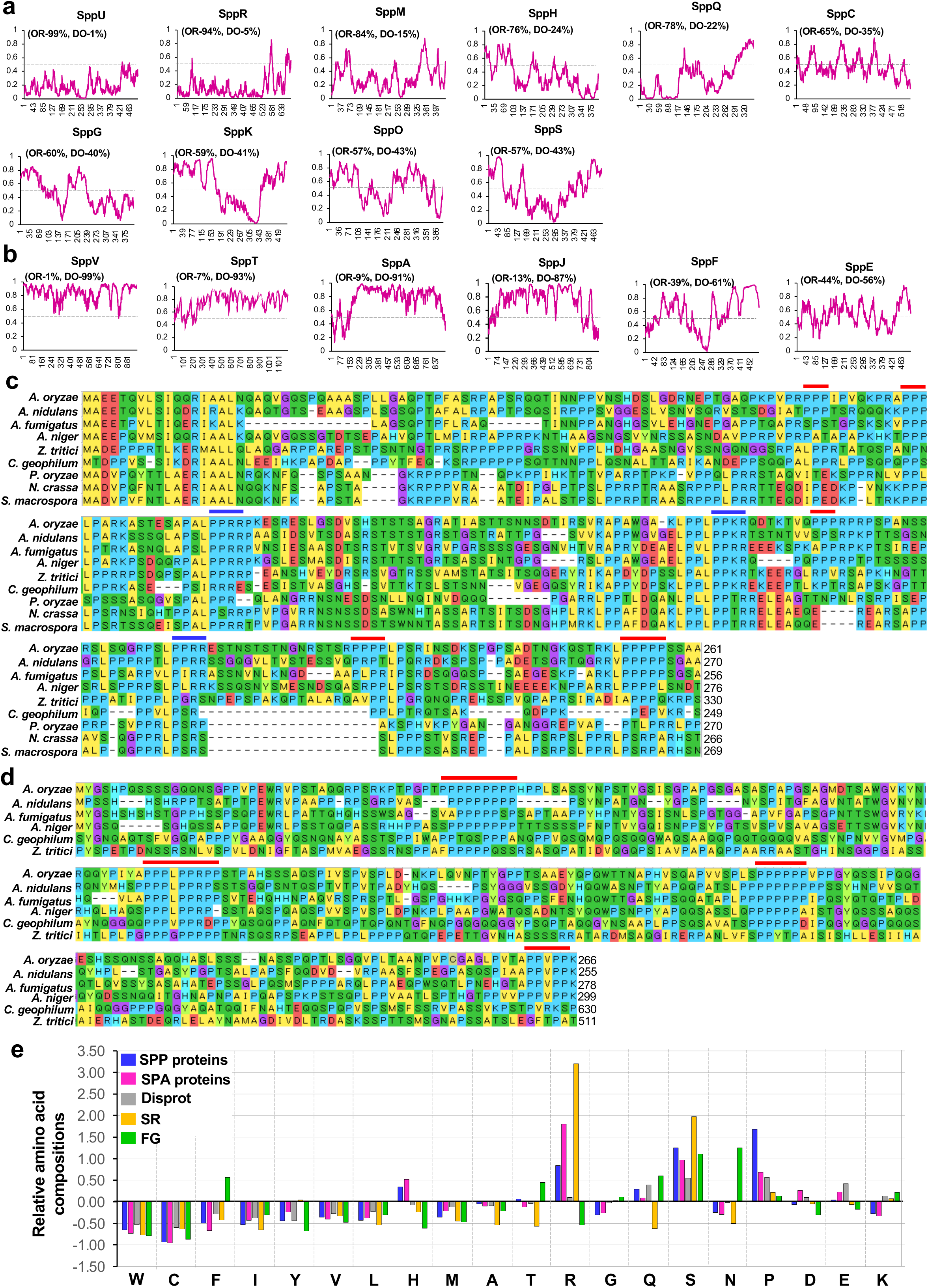
Prediction of disordered regions of SPP proteins. **a** SPP proteins containing a higher portion of the ordered region. **b** SPP proteins containing a higher portion of the disordered region. In the graph of prediction of disordered regions with IUPred2A, they axis indicates the predicted probability of disorder, and the x-axis represents the amino acid sequence. OR; ordered, DO; disordered. **c and d** Multiple sequence alignment of the N-terminal disordered regions of SppB and SppN. Amino acid sequences of orthologous proteins were retrieved from NCB!using protein BLAST. Multiple sequence alignment was made using ClustaiW in MegaX. Red lines upper to the sequence denote conserved poly-proline motifs. Blue lines denote conserved PPRRIPPKR motifs. e Calculation of relative amino acid composition in the disordered region of SPP and comparison with the disordered regions of SPA, SR FG, and DisProt datasets.

**Supplementary Figure 8:**
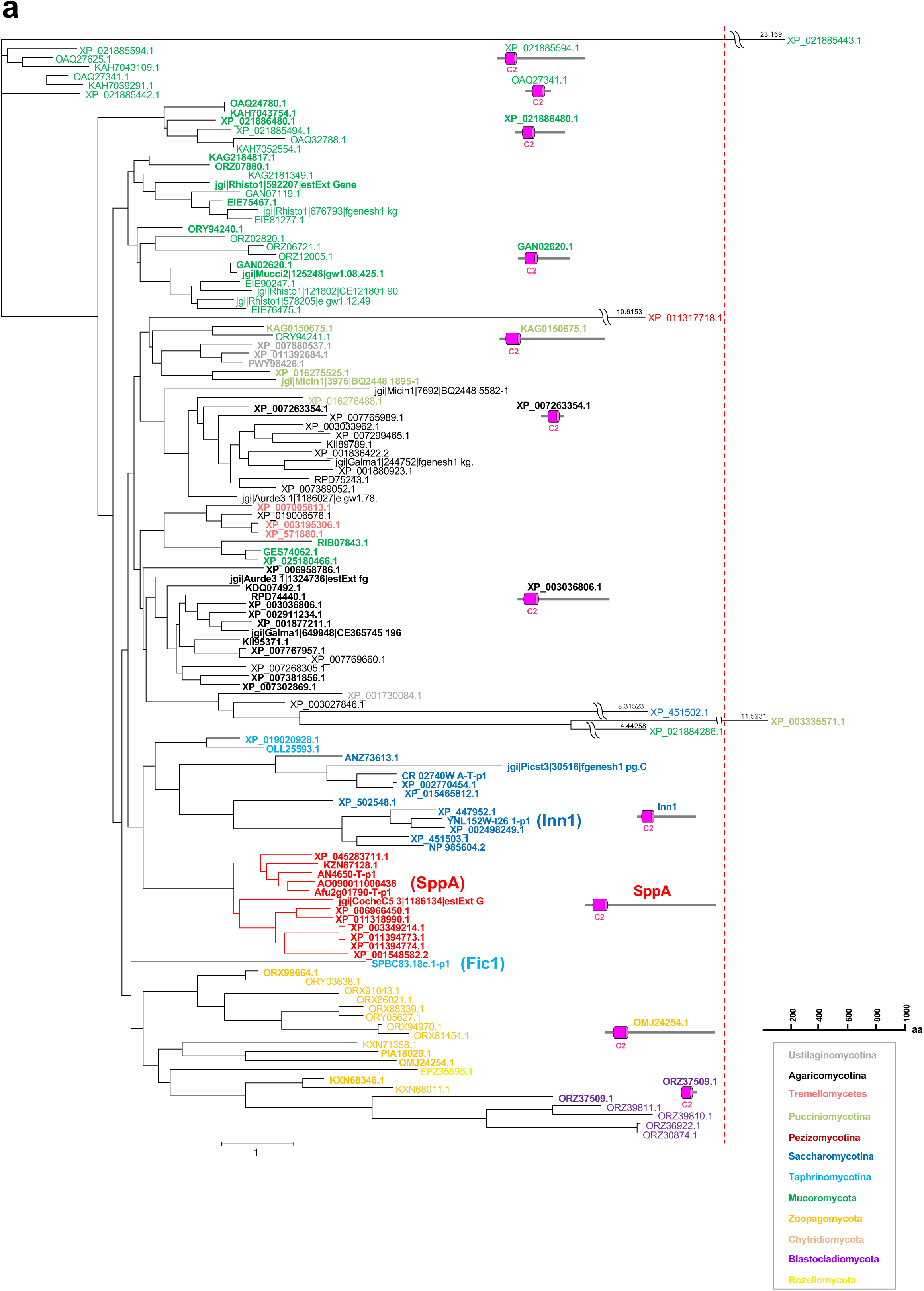
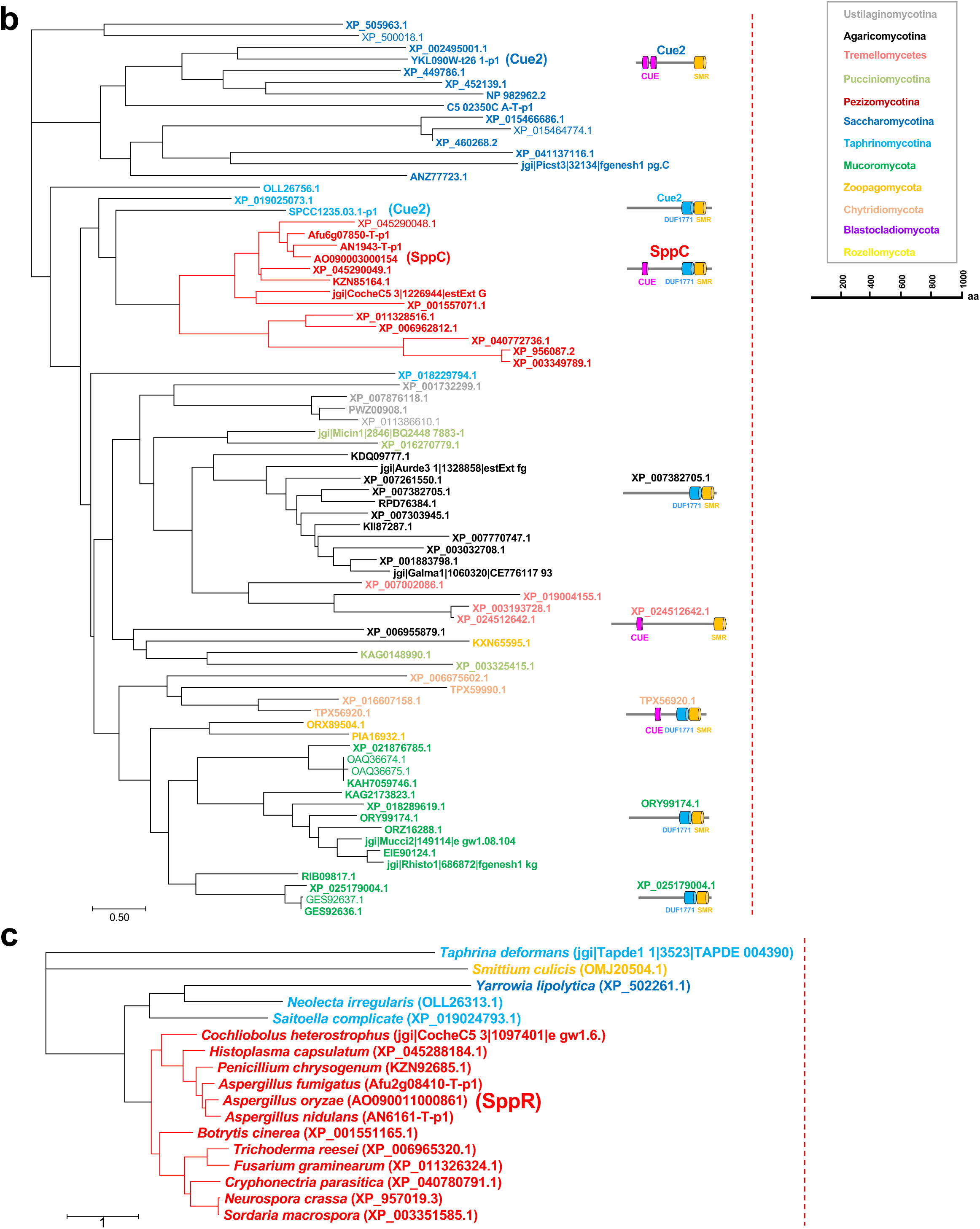
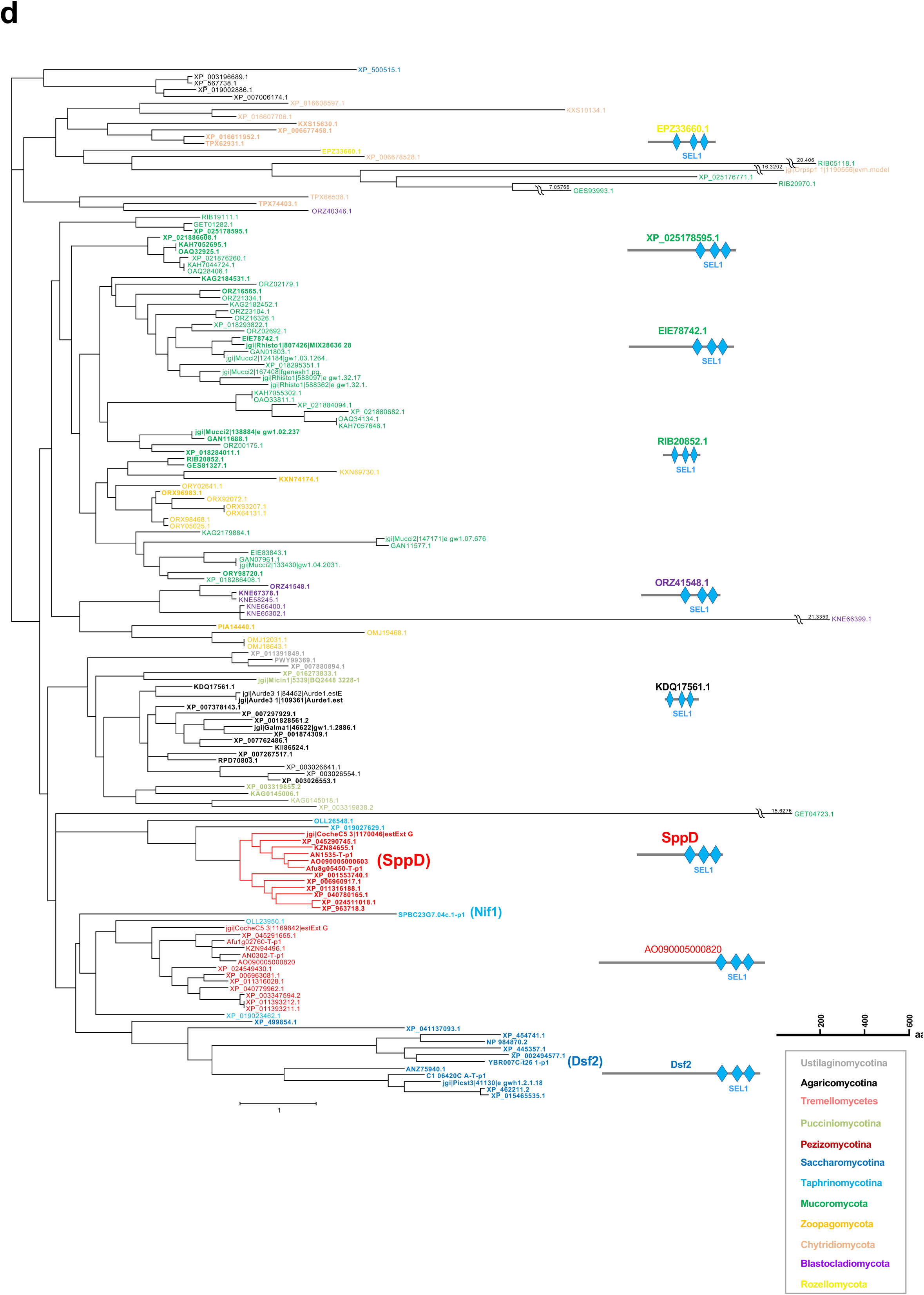
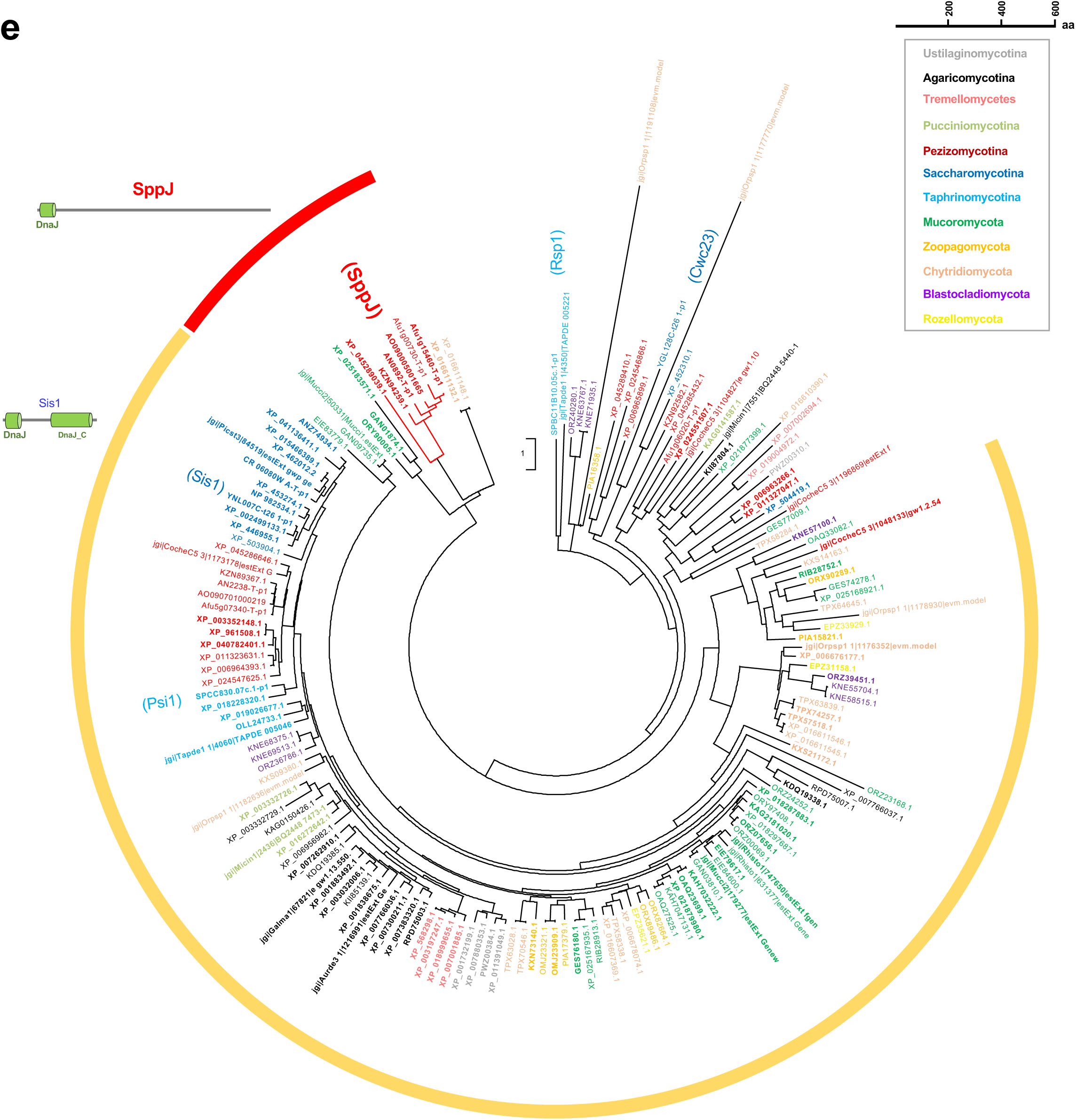
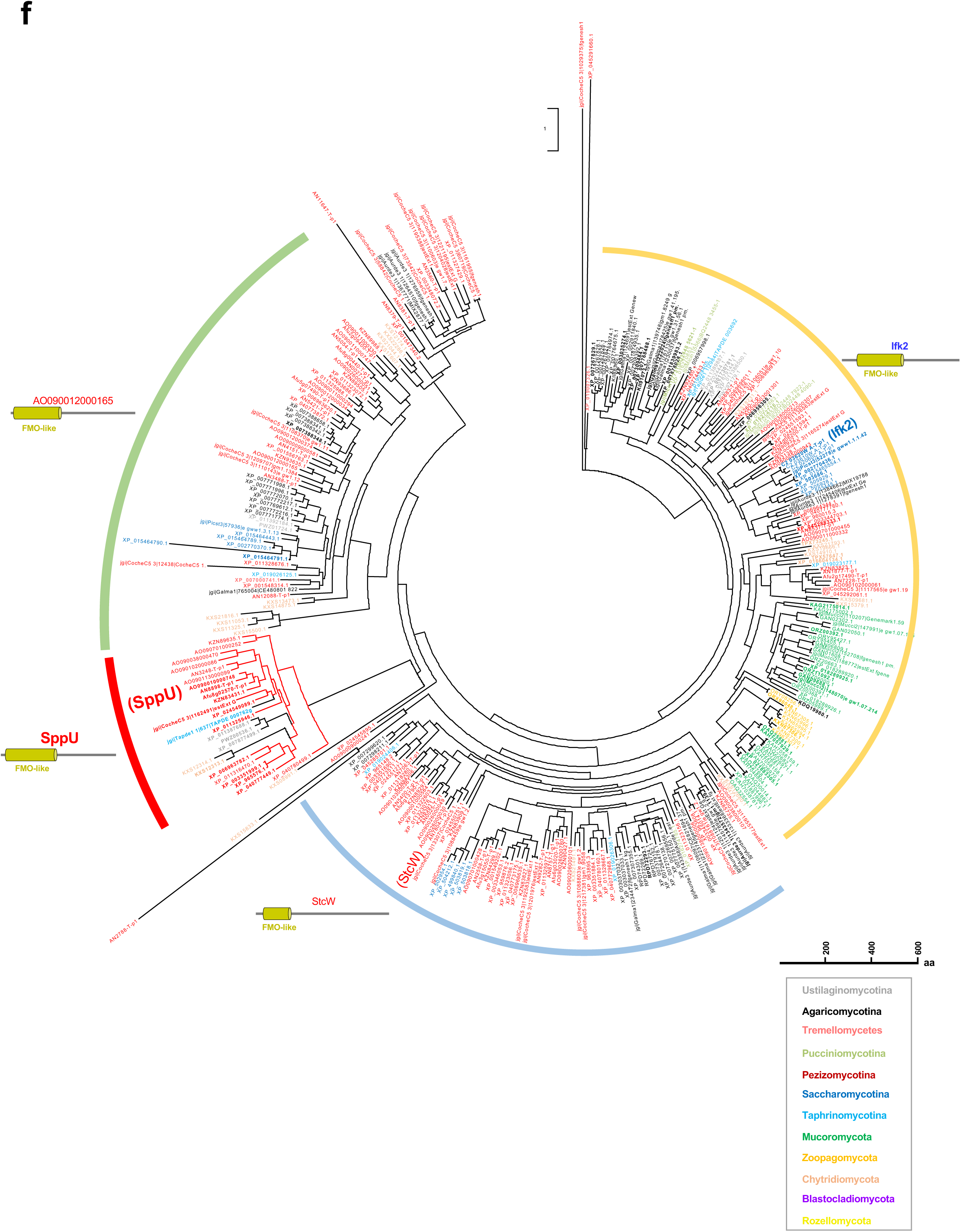
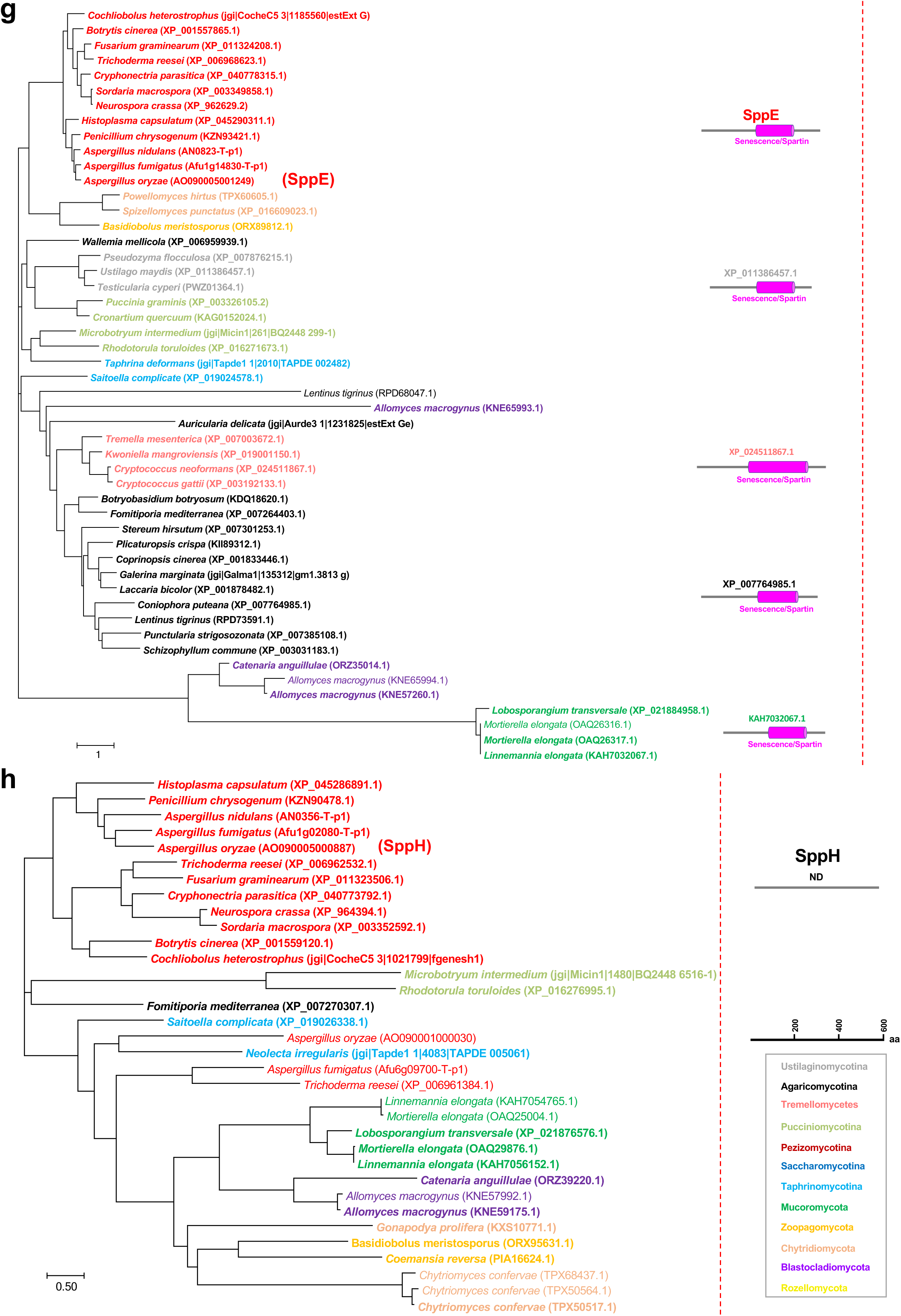
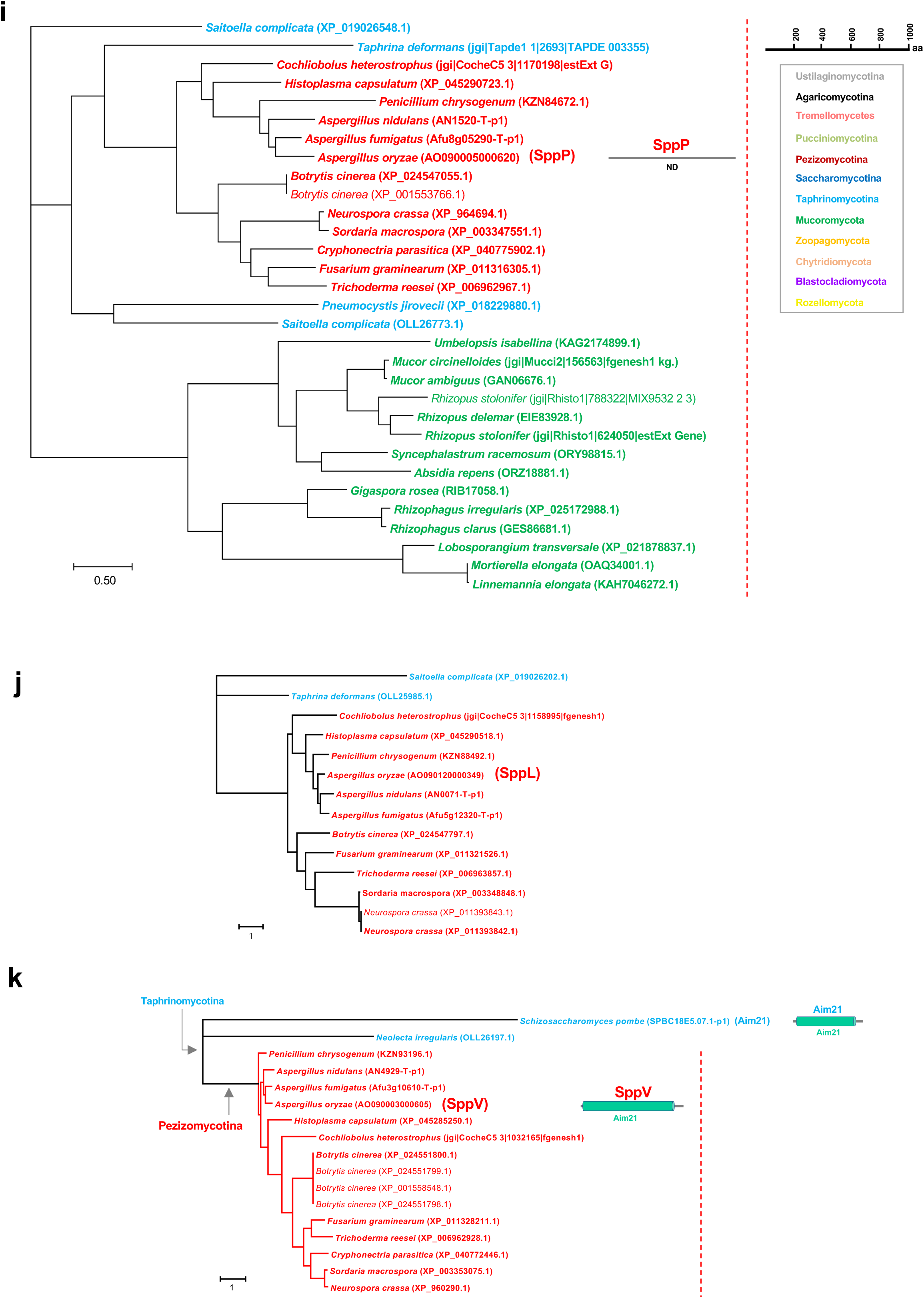
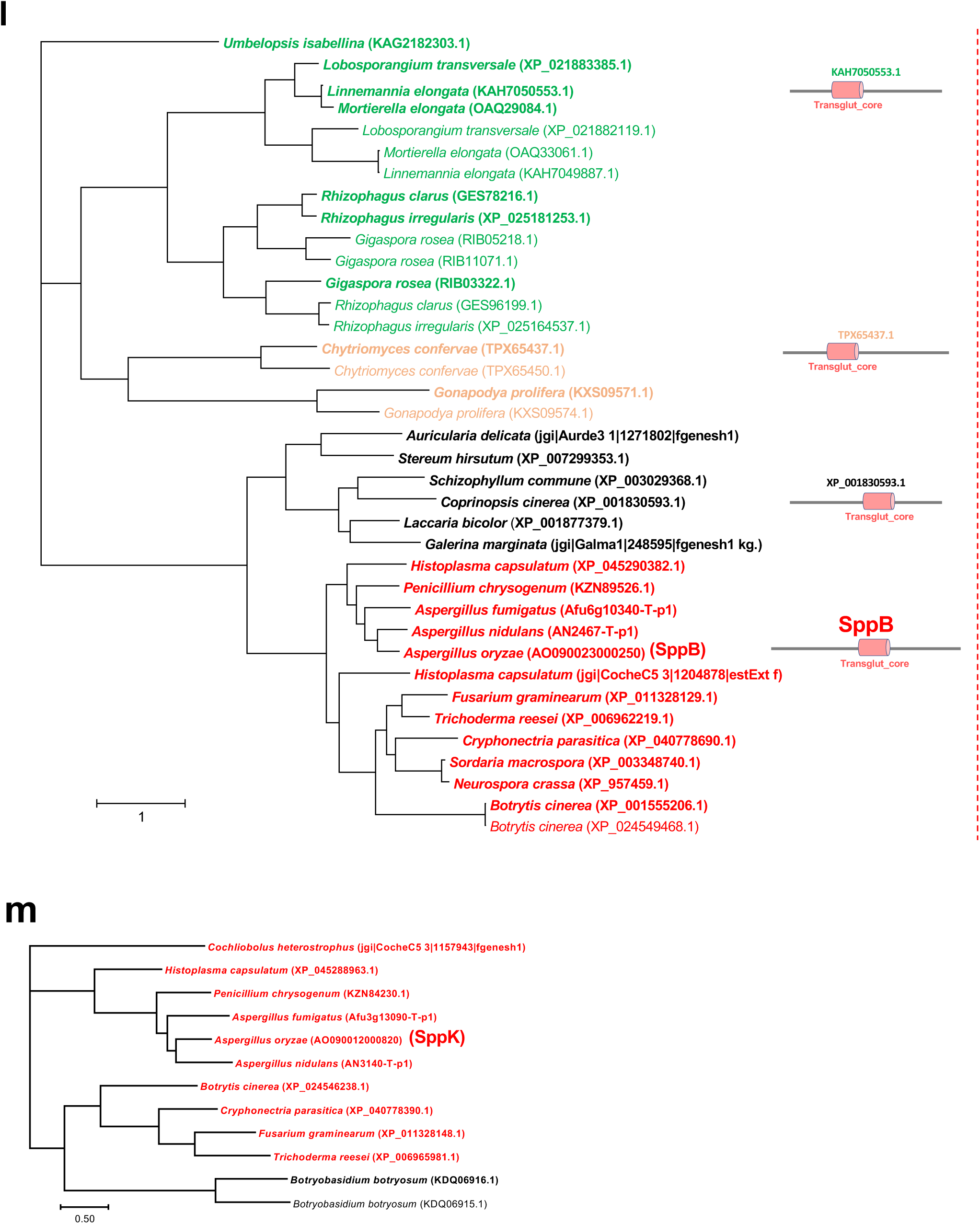
Phylogenies of 13 SPP proteins having orthologs outside of Pezizomycotina. Maximum-likelihood trees were made using Mega 7 for proteins belonging to the same orthologous groups classified by OrthoFinder. Domains were predicted using the Simple modular architecture research tool (SMART). Diagrams of protein domain structures are shown with the respective clades. Bold letters represent proteins used for the substitution rate analysis in Fig. 7. **a-m** Phylogenies of proteins in the orthologous groups including SPP proteins.

**Supplementary Figure 9:**
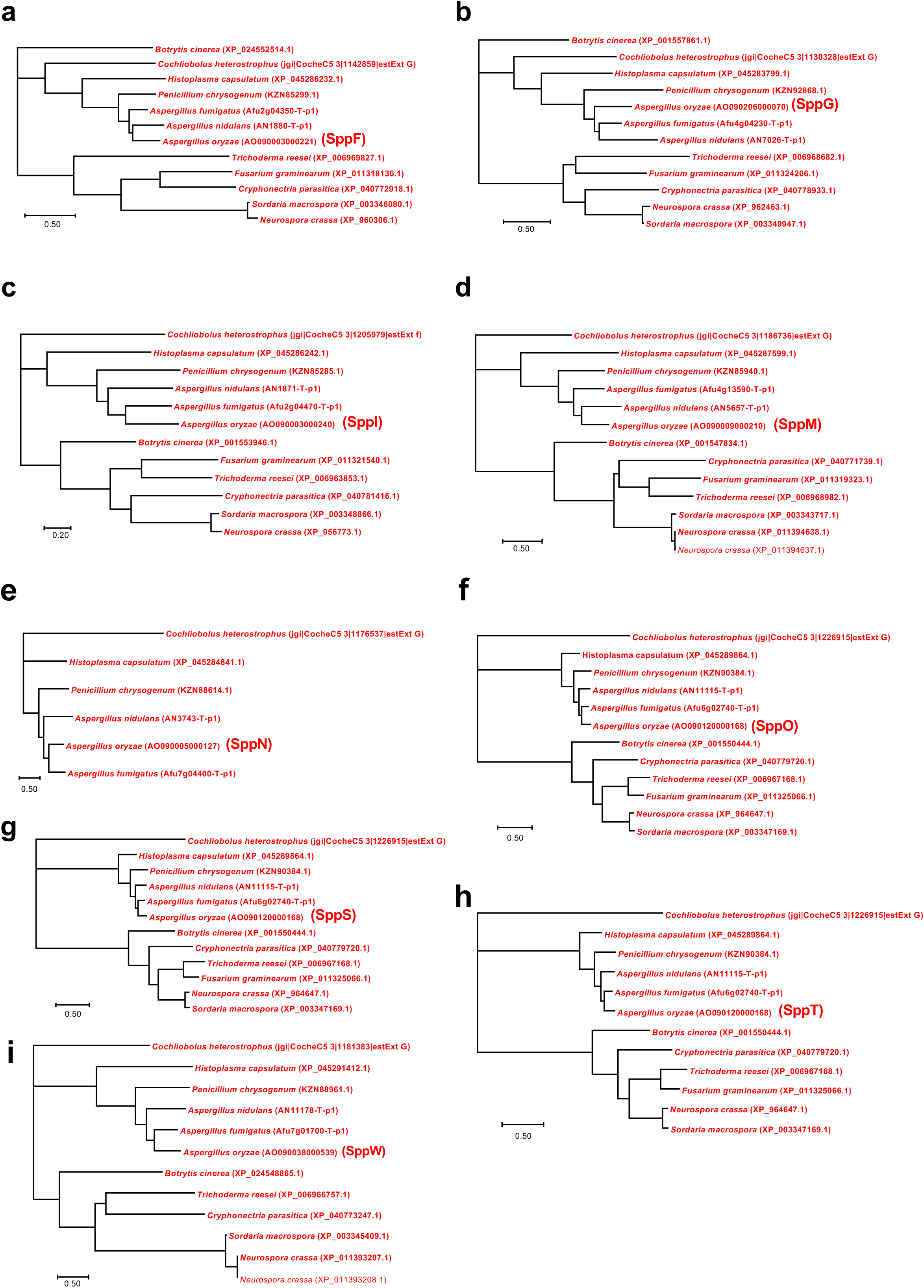
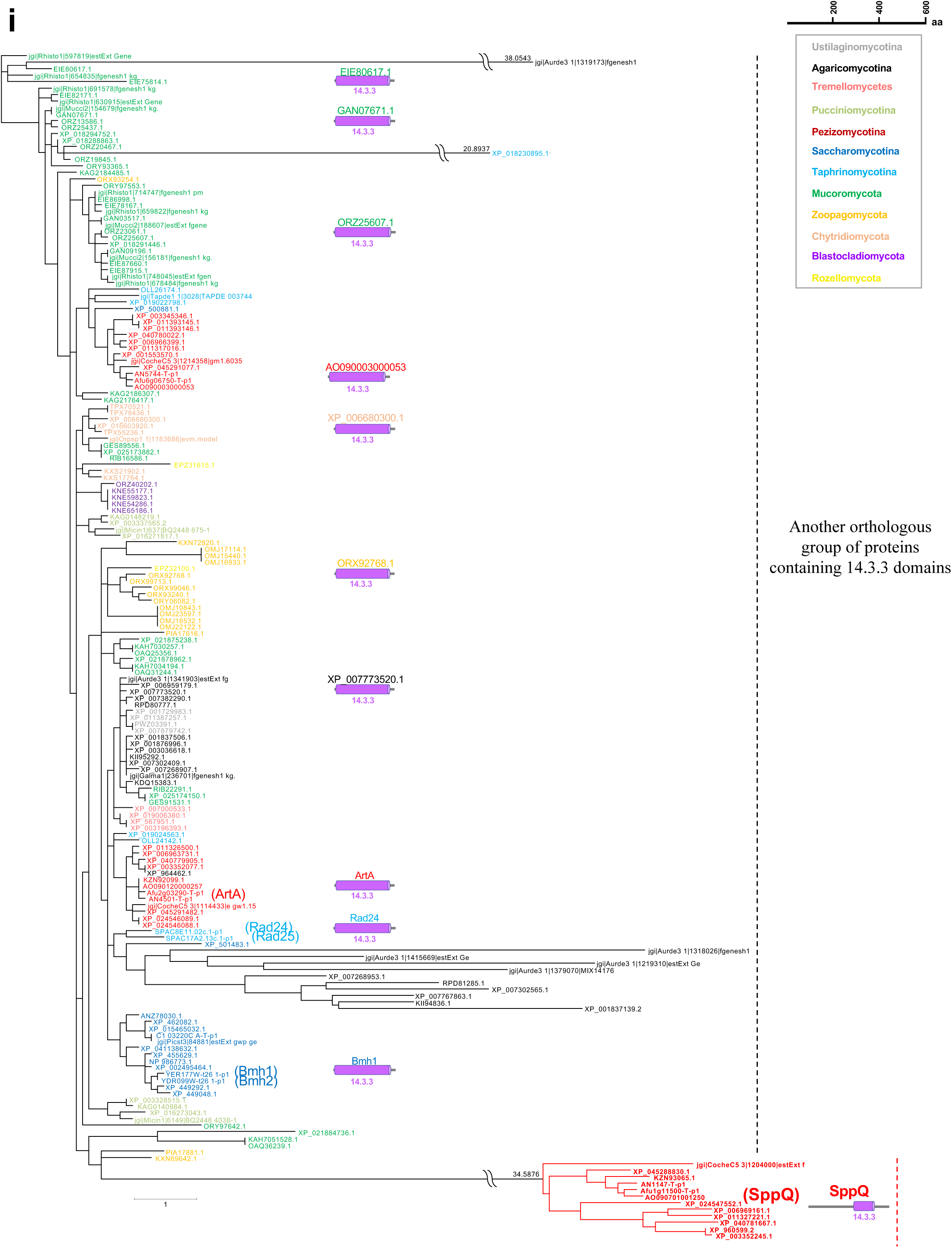
Phylogenies of Pezizomycotina-specific SPP proteins. Maximum-likelihood trees were made using Mega 7 for proteins belonging to the orthologous groups classified by OrthoFinder. Domains were predicted using the Simple modular architecture research tool (SMART). Diagrams of protein domain structures are shown with the respective clades. Bold letters represent the proteins used for substitution rate analysis in Fig. 7. **a-h** Phylogenies of proteins in the same orthologous groups including SPP proteins. **i** Phylogeny of proteins in the orthologous group including SppQ. Note that the phylogeny also includes another orthologous group proteins containing 14.3.3 domains.

**Supplementary Figure 10:**
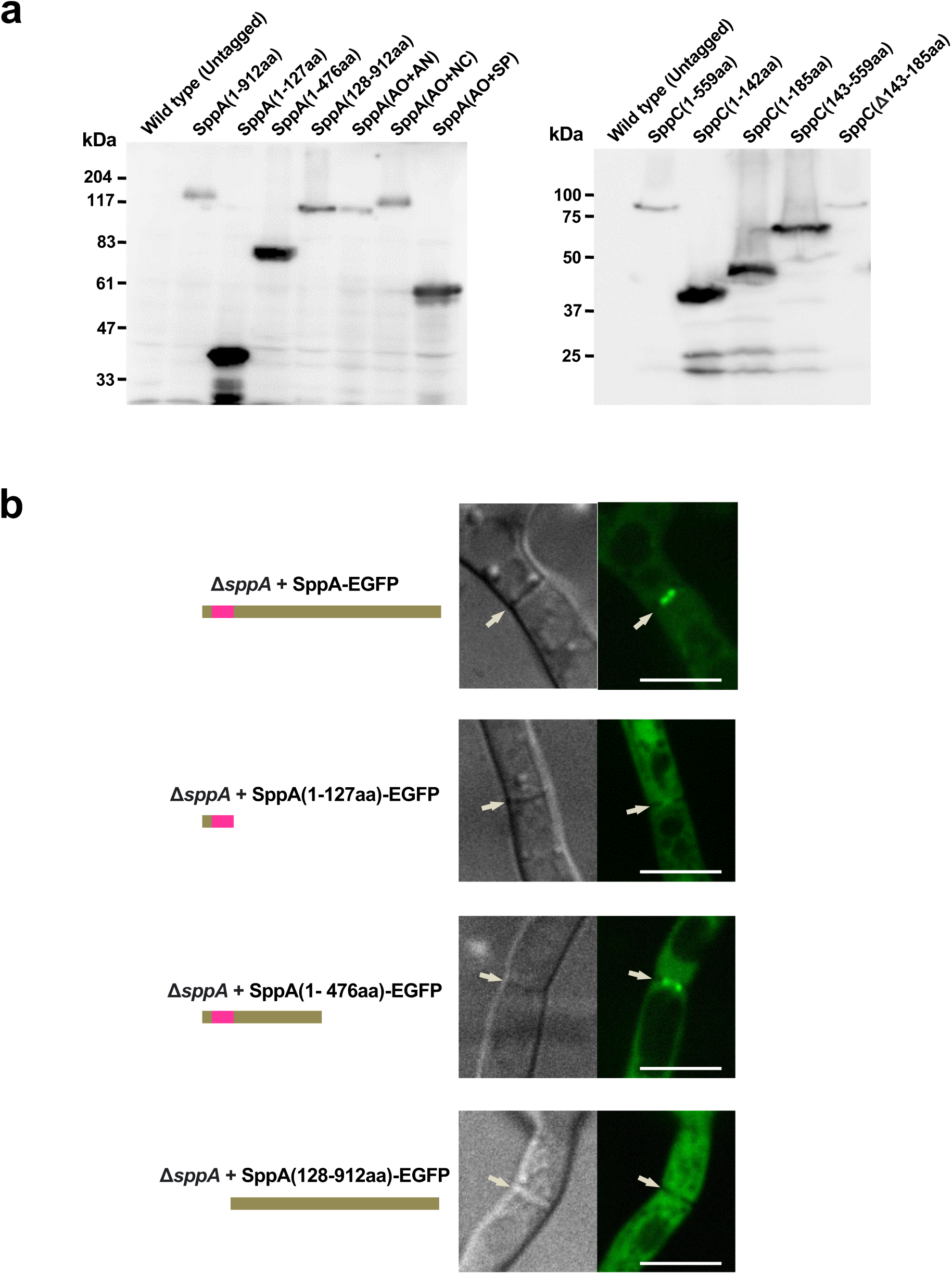
Expression of SppA and SppC truncations. **a** Expression of SppA/SppC variants, including truncations and chimeric proteins, confirmed by western blotting. Crude proteins (20 µg) isolated from the individual strain grown in DPY liquid medium were loaded in each lane. Expression of SppA/C and truncated/chimeric variants fused with EGFP was confirmed by Western blotting with anti-GFP antibody. Note that low-molecular weight variants showed high signal intensities due to high transfer efficiency. **b** The N-terminus of SppA is essential for its localization around the septal pore. Single confocal fluorescence images are shown, and arrows indicate apical septa. Scale bars, 5 μm.

**Supplementary Figure 11:**
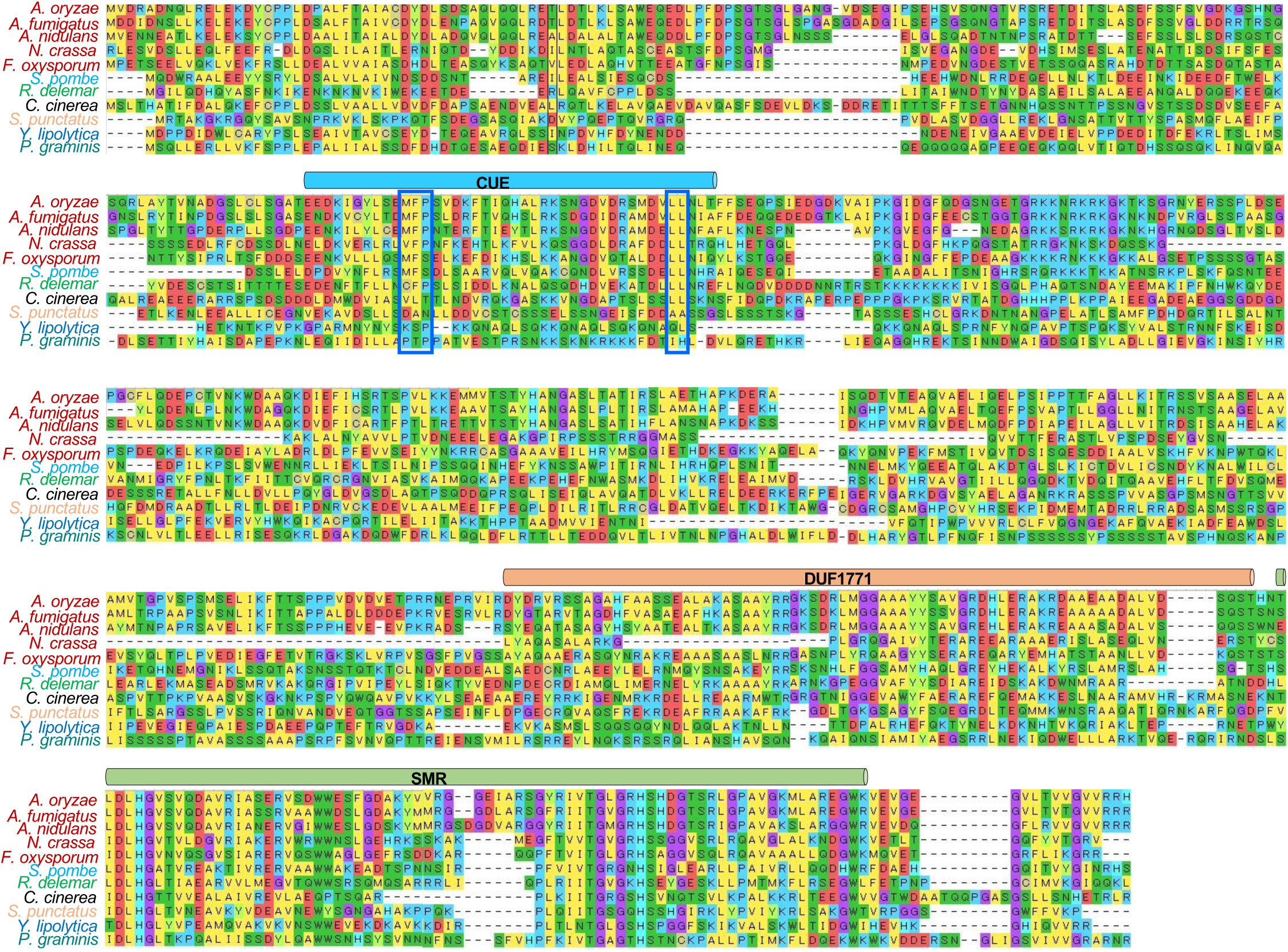
Multiple sequence alignment of SppC and orthologs. Blue boxes represent two conserved motifs (MFP and LL) essential for ubiquitin-related function.

## Notes

### Competing Interest Statement

The authors have declared no competing interest.

### Summary of Updates

In the main body, I have found the shadow background, which I have corrected. Additionally, I have corrected the order of authors appearing on the site.

